# Nucleosomal Asymmetry Shapes Histone Mark Binding and Promotes Poising at Bivalent Domains

**DOI:** 10.1101/2021.02.08.430127

**Authors:** Elana Bryan, Marie Warburton, Kimberly M. Webb, Katy A. McLaughlin, Christos Spanos, Christina Ambrosi, Viktoria Major, Tuncay Baubec, Juri Rappsilber, Philipp Voigt

## Abstract

Promoters of developmental genes in embryonic stem cells (ESCs) are marked by histone H3 lysine 4 trimethylation (H3K4me3) and H3K27me3 in an asymmetric nucleosomal conformation, with each sister histone H3 carrying only one mark. These bivalent domains are thought to poise genes for timely activation upon differentiation. Here we show that asymmetric bivalent nucleosomes recruit repressive H3K27me3 binders but fail to enrich activating H3K4me3 binders, despite presence of H3K4me3, thereby promoting a poised state. Strikingly, the bivalent mark combination further attracts chromatin proteins that are not recruited by each mark individually, including the histone acetyltransferase complex KAT6B (MORF). Knockout of KAT6B blocks neuronal differentiation, demonstrating that bivalency-specific readers are critical for proper ESC differentiation. These findings reveal how histone mark bivalency directly promotes establishment of a poised state at developmental genes, while highlighting how nucleosomal asymmetry is critical for histone mark readout and function.

## Introduction

Histone modifications have emerged as key regulators of transcription and other chromatin-templated processes. Often in combinatorial fashion, histone marks set up chromatin environments that reflect, reinforce, and potentially instruct transcriptional states by directly regulating access to the DNA or by recruiting factors that activate or repress transcription. Promoters of developmental genes in embryonic stem cells (ESCs) are marked by a distinctive histone modification signature comprised of the active histone mark histone H3 lysine 4 trimethylation (H3K4me3) and the repressive mark H3K27me3 (Azuara et al., 2006; Bernstein et al., 2006; Harikumar and Meshorer, 2015; Mikkelsen et al., 2007; Voigt et al., 2013). These so-called ‘bivalent domains’ are presumed to poise expression of developmental genes primarily in ESCs by maintaining a repressed but plastic state that allows for prompt activation or stable repression of these genes upon differentiation cues. Bivalent domains are established by the histone methyltransferase complexes MLL2 and Polycomb repressive complex (PRC) 2, catalyzing H3K4me3 and H3K27me3, respectively (Christophersen and Helin, 2010; Piunti and Shilatifard, 2016; Schuettengruber et al., 2017; Yu et al., 2019). Supporting a pivotal role of bivalent domains in regulating developmental genes, knockouts or loss-of-function mutations in these complexes abolish bivalency and lead to developmental defects in mice and compromised differentiation potential in ESCs (Laugesen and Helin, 2014; Piunti and Shilatifard, 2016; Vastenhouw and Schier, 2012). However, the mechanisms by which bivalent domains poise genes for expression are largely unclear and it remains elusive whether the bivalent histone marks themselves are key drivers of poising.

Given the central role of binding proteins or ‘readers’ in executing histone mark function, proteins that bind H3K4me3 or H3K27me3 are candidates for translating bivalency into a poised state via a histone mark-based mechanism. Complexes featuring binding proteins for the active H3K4me3 mark comprise the general transcription factor TFIID, transcriptional co-activators, histone modifiers, and chromatin remodelers, whereas the repressive H3K27me3 mark is primarily recognized by PRC2 and canonical PRC1 complexes (Bartke et al., 2010; Eberl et al., 2013; Musselman et al., 2012; Taverna et al., 2007; Vermeulen et al., 2010). However, thus far recognition of H3K4me3 and H3K27me3 has only been studied individually, rendering potential antagonistic or multivalent synergistic effects of their bivalent coexistence elusive. Moreover, we and others have shown that bivalent nucleosomes are asymmetrically modified, carrying H3K4me3 and H3K27me3 on two separate histone H3 copies, rather than featuring two histone H3 copies modified with both H3K4me3 and H3K27me3 (Shema et al., 2016; Voigt et al., 2012). However, it remains unclear how nucleosomal asymmetry affects recruitment of histone mark binding proteins—and thus histone mark function—at bivalent nucleosomes and beyond.

Here, we set out to determine how asymmetric bivalent nucleosomes are decoded in order to clarify whether the bivalent marks directly act to poise developmental genes in ESCs. Using an approach based on nucleosome pulldown assays with recombinant, site-specifically modified nucleosomes followed by quantitative mass spectrometry (MS) analysis, we reveal that bivalent nucleosomes recruit repressive complexes as well as bivalency-specific binders, but fail to recruit H3K4me3 binders. These findings provide evidence for a histone mark-based poising mechanism and illustrate how nucleosomal asymmetry regulates histone mark function.

## Results

### Monovalent asymmetric nucleosomes fail to recruit H3K4me3 and H3K27me3 binding proteins

In order to resolve how the asymmetric bivalent mark combination might directly establish or facilitate poising at bivalent domains, we generated site-specifically modified histones via native chemical ligation and assembled them into recombinant nucleosomes carrying symmetric and asymmetric H3K4me3 and H3K27me3 (Figure S1). We then carried out nucleosome pulldown assays with mouse E14 ESC nuclear extract (NE) followed by label-free quantification (LFQ) liquid chromatography (LC)-coupled mass spectrometry (MS) analysis (Figure 1A). We first sought to determine how asymmetric monovalent presence of H3K4me3—as observed at bivalent domains *in vivo—*regulates its readout and affects recruitment of H3K4me3 binding proteins. As expected from previous studies (Bartke et al., 2010; Eberl et al., 2013), when performing pulldowns with symmetric H3K4me3 (denoted as H3K4me3/3) nucleosomes, we observed specific enrichment of a host of known H3K4me3 binding proteins, including members of the TFIID, SETD1, and SIN3A/B complexes, chromatin remodelers such as NURF, CHD1, and CHD8, and other PHD finger proteins such as PHF2 (Figures 1B and 1C, Table S1). We further observed depletion of HMG20B, PHF21A, G9a/GLP, and PRC2 on H3K4me3/3 nucleosomes (Figures 1B and 1C, Table S1). Strikingly, when we performed analogous pulldown experiments with asymmetric (‘H3K4me0/3’) nucleosomes, enrichment of H3K4me3 binders and depletion of repressive factors was lost (Figures 1B and 1C, Table S1), indicating that asymmetric nucleosomes fail to recruit H3K4me3 binding proteins, despite presence of the mark on one of the two histone H3 copies in the nucleosome. In line with these findings, asymmetric H3K27me3 nucleosomes likewise failed to enrich known H3K27me3 binding proteins, while symmetric H3K27me3 nucleosomes robustly recruited known H3K27me3 binders (Figures 1B and 1D, Table S2). Taken together, symmetric modification appears to be required to achieve enrichment of binding proteins for both H3K4me3 and H3K27me3 in nucleosome pulldown assays, while recruitment was diminished for asymmetric nucleosomes. These findings suggest that loci featuring asymmetric nucleosomes—such as bivalent domains—are functionally distinct from loci containing nucleosomes symmetrically modified with H3K4me3 or H3K27me3.

**Figure 1.**
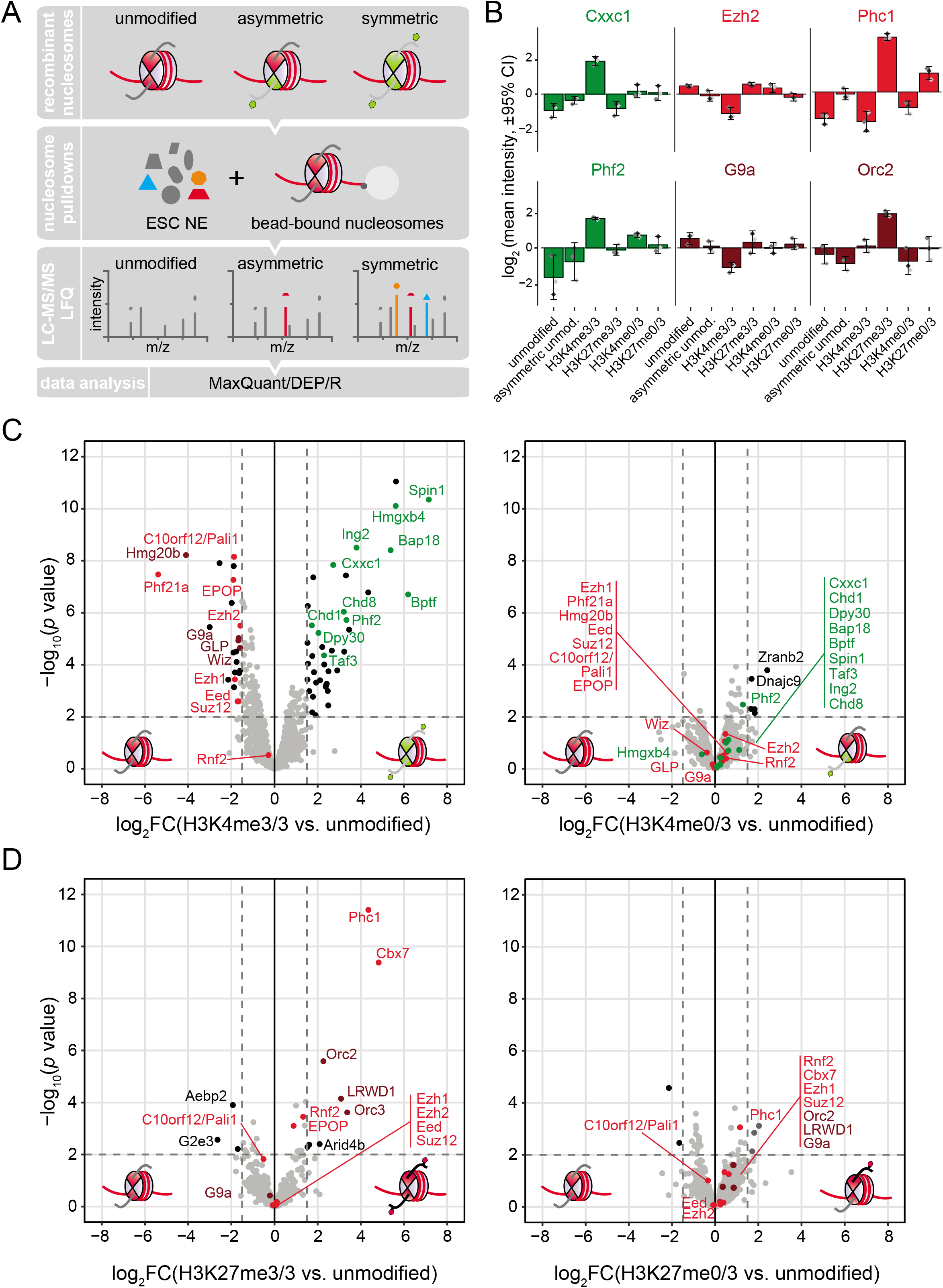
Monovalent asymmetric nucleosomes fail to recruit H3K4me3 and H3K27me3 binding proteins. **(A)** Schematic overview of the experimental approach used to determine the impact of asymmetric modification on histone mark binding. **(B)** Enrichment and depletion of representative proteins on nucleosomes symmetrically or asymmetrically modified by H3K4me3 or H3K27me3. Plotted are label-free quantification (LFQ) intensities expressed as fold change relative to the mean LFQ intensity of all pulldown conditions analyzed. Error bars represent 95% confidence intervals of means, and diamonds denote individual values of three independent experiments. Green bars, H3K4me3 binding proteins and associated complex members; red bars, H3K27me3 binding proteins and associated PRC members; dark red bars, non-PRC H3K27me3 binding proteins and associated factors; H3K4me3/3, symmetric modification with H3K4me3; H3K4me0/3, asymmetric modification with H3K4me3 on one copy of histone H3 per nucleosome. Analogous notations used for H3K27me3-modified nucleosomes. **(C,D)** Volcano plots of LFQ MS analysis of nucleosome pulldown experiments with symmetric (left panels) and asymmetric (right panels) nucleosomes modified with H3K4me3 (C) and H3K27me3 (D), compared against unmodified symmetric and asymmetric (containing C-terminally tagged histone H3) controls. The C-terminal tags present on histone H3 in asymmetric nucleosomes did not significantly affect binding of the proteins studied here (see Extended Data Fig. 1c). Means and *p* values are derived from three independent pulldown experiments. Black dots, significantly enriched or depleted proteins (log2 fold change ≤ 1.5 or ≥ 1.5, respectively, and *p* value ≤ 0.01), color scheme for known H3K4me3 and H3K27me3 binders as in (B). See also Figure S1 and Tables S1 and S2.

### Symmetric modification enhances the apparent affinity of H3K4me3 and H3K27me3 binders towards nucleosomes

We next sought to clarify why recruitment of mark binding proteins was markedly less efficient for asymmetric compared to symmetric nucleosomes, despite carrying the same modifications. To this end, we performed a series of nucleosome pulldown assays with symmetric and asymmetric nucleosomes and analyzed recruitment of select H3K4me3 binders in a semi-quantitative fashion using titration experiments (Figures 2A and S2A). Robust, concentration-dependent binding of TAF3, PHF2, and ING2 was detected for symmetric H3K4me3 nucleosomes, whereas binding to asymmetric H3K4me3 nucleosomes was substantially weaker (Figure 2A), in line with our LFQ-MS analysis (see Figure 1B and 1C). We reasoned that the marked decrease in apparent affinity towards asymmetric nucleosomes could be due to factors binding to the unmodified tail, precluding recruitment of H3K4me3 binders due to steric exclusion. However, when performing pulldowns with asymmetric nucleosomes containing a tail-less rather than full-length histone H3 as the unmodified counterpart, enrichment of H3K4me3 binders was not recovered (Figure 2A). We next hypothesized that the two-fold higher abundance of binding sites on symmetric nucleosomes enhances interaction with binding proteins based on mass action. However, while partially enhancing ING2 binding, matching the abundance of H3K4me3 sites by doubling the amount of asymmetric nucleosomes did not recover binding of TAF3 and PHF2 to levels seen for symmetric nucleosomes (Figure 2A). These findings indicate that the symmetric arrangement itself—rather than mere increased overall mark abundance—favors recruitment of H3K4me3 binders to symmetric nucleosomes.

**Figure 2.**
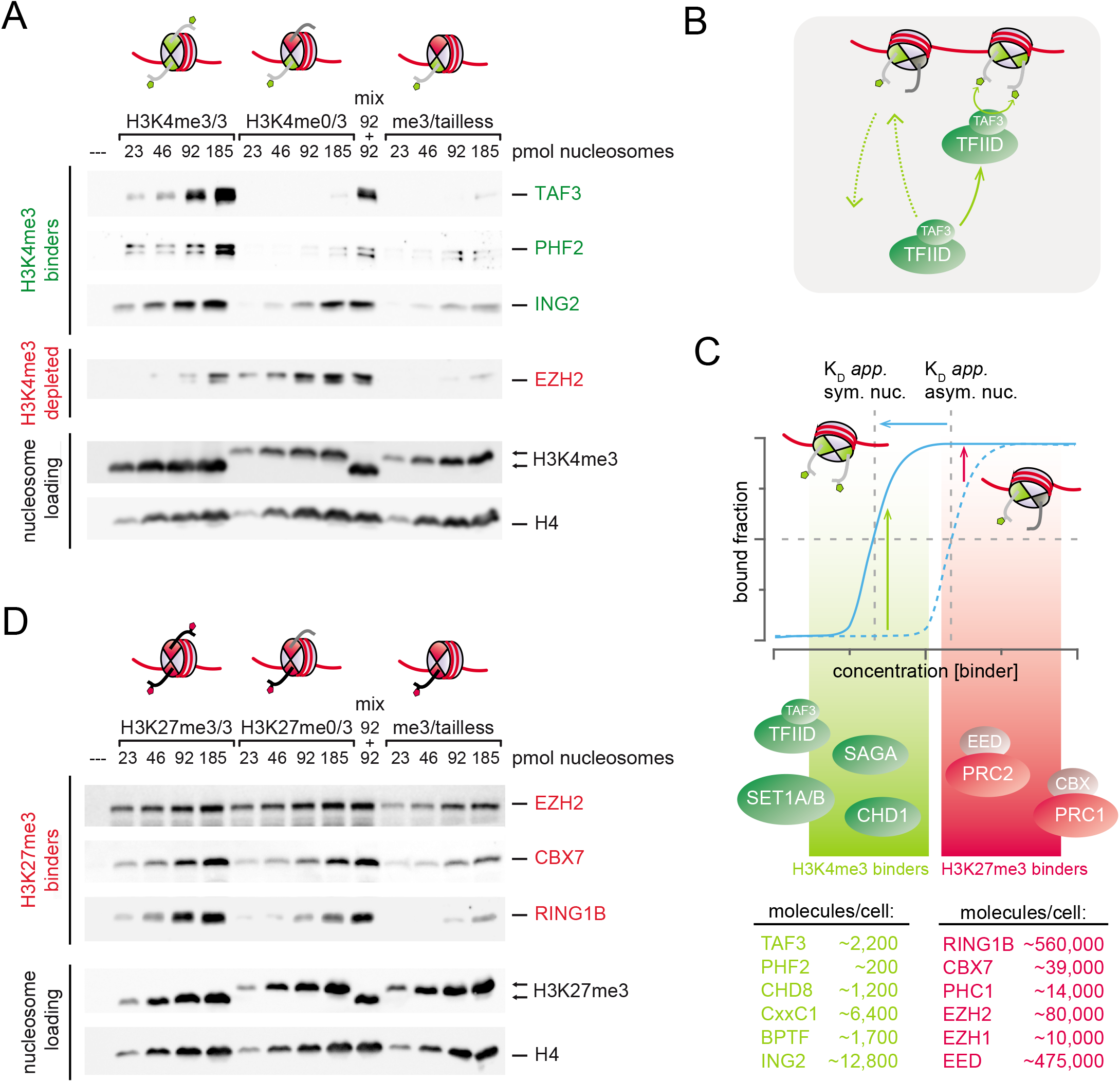
Symmetric modification enhances the apparent affinity of H3K4me3 and H3K27me3 binders towards nucleosomes. **(A)** Western Blot analysis of titration pulldown assays with nucleosomes carrying H3K4me3 in symmetric, asymmetric, and asymmetric-tailless conformation. The pulldown labelled ‘mix’ was performed with an equimolar mixture of symmetric and asymmetric nucleosomes. Blots shown are representative of three independent experiments. **(B)** Scheme illustrating how allovalent effects increase apparent affinity towards symmetric nucleosomes. Availability of two binding sites enhances recruitment while concomitantly allowing for rebinding to the neighboring mark upon dissociation, retaining the binder on the nucleosome. **(C)** Allovalency shifts the binding equilibrium towards the bound state for symmetric nucleosomes. The resulting gain in binding is expected to be stronger for factors with concentrations below the apparent K_D_ compared to highly abundant binders, which would experience only minor gains in binding. Intensity-based absolute quantification (iBAQ)-derived protein copy numbers per cell for representative H3K4me3 and H3K27me3 binders are from Zhang et al. (2017). **(D)** Western Blot analysis of titration pulldown assays with nucleosomes carrying H3K27me3 in symmetric, asymmetric, and asymmetric-tailless conformation. Blots shown are representative of three independent experiments. See also Figure S2.

On symmetric nucleosomes, two equivalent binding sites are available for reader proteins that contain a single cognate binding domain. This situation is akin to clusters of near-equivalent phosphorylation sites that recruit signaling factors containing a single phosphorylation binding domain. Conceptually distinct from multivalency, the term ‘allovalency’ has been coined to describe the behavior of such systems that consist of a single binding domain interacting with a polyvalent ligand containing multiple identical binding sites in tandem (Ivarsson and Jemth, 2019; Klein et al., 2003; Locasale, 2008; Mittag et al., 2008; Olsen et al., 2017). Upon dissociation from one binding site, the binder can rapidly rebind another epitope nearby before leaving the ‘capture sphere’ of the polyvalent ligand, resulting in rapid intracomplex exchange rather than complete dissociation, thus increasing overall apparent affinity (illustrated in Figure 2B for TAF3). As observed in our nucleosome pulldown assays, these effects increase the apparent affinity of binders towards symmetric H3K4me3 modification, resulting in markedly enhanced binding (Figure 2A, illustrated in Figure 2C).

Increased apparent affinity through allovalent effects on symmetric nucleosomes and the associated shift in apparent K_D_ would be expected to lead to larger gains in binding for lowly compared to highly abundant binders that are already approaching saturated nucleosome binding (illustrated in Figure 2C). Interestingly, most H3K4me3 binders are expressed at low levels in ESCs, whereas PRC1 and PRC2 are considerably more abundant (Zhang et al., 2017). Indeed, while the lowly abundant H3K4me3 binders exhibited markedly increased binding to symmetric nucleosomes, this increase was much less pronounced for H3K27me3 binders (Figures 2D and S2B). These findings indicate that the high abundance of PRC1 and PRC2 complexes in ESCs overcomes the requirement of symmetric modification for efficient recruitment, suggesting that nucleosomal asymmetry at bivalent nucleosomes in ESCs would impact recruitment of H3K4me3 binders more strongly than H3K27me3 binders.

### Bivalent nucleosomes recruit repressive H3K27me3 binders and bivalency-specific binders, but not H3K4me3 binders

Having established that nucleosomal asymmetry affects the readout of monovalent H3K4me3 and, to a lesser extent, H3K27me3, we next sought to determine whether these effects are recapitulated for asymmetric bivalent nucleosomes by performing pulldown experiments with bivalent nucleosomes. In line with the different sensitivity of H3K4me3 and H3K27me3 reader binding towards asymmetric modification (Figures 1 and 2), asymmetric bivalent nucleosomes failed to recruit H3K4me3 binding proteins such as TAF3, PHF2, and ING2, but significantly enriched H3K27me3 binders (Figures 3A–3C and S3A, Table S3). In the case of PRC1, enrichment on bivalent nucleosomes was lower compared to symmetric H3K27me3 nucleosomes (Figures 3B and S3A), as expected from the reduced affinity of H3K27me3 binders towards asymmetric H3K27me3 nucleosomes (Figures 1D and 2D). Interestingly, along with the core subunit EZH2, the auxiliary PRC2 subunits EPOP and EZHIP were specifically enriched on bivalent nucleosomes (Figures 3B and S3A), suggesting differential interaction of specific PRC2 subcomplexes with bivalent nucleosomes.

**Figure 3.**
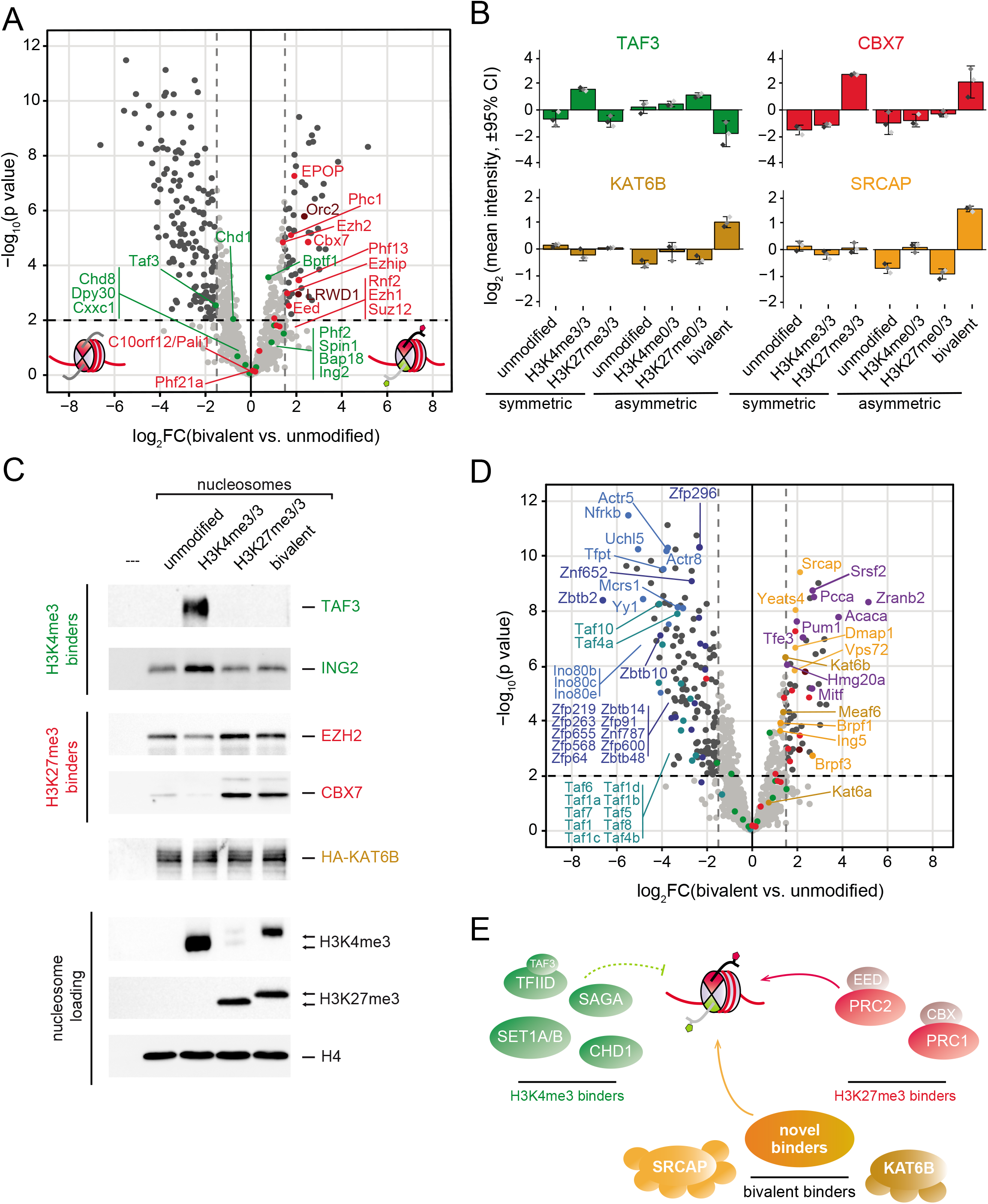
Bivalent nucleosomes recruit repressive H3K27me3 binders and bivalency-specific binders, but not H3K4me3 binders. **(A)** Volcano plot analysis of LFQ MS data for nucleosome pulldown experiments with bivalent nucleosomes compared against an unmodified asymmetric (containing C-terminally tagged histone H3) nucleosome control. Means and *p* values are derived from three independent pulldown experiments. Dark gray dots, significantly enriched or depleted proteins (log2 fold change ≤ 1.5 or ≥ 1.5, respectively, and *p* value ≤ 0.01). Known H3K4me3 and H3K27me3 binders are highlighted by color as in Figure 1. **(B)** Enrichment and depletion of representative proteins on symmetric as well as monovalent and bivalent asymmetric nucleosomes. Plotted are fold changes of LFQ intensities relative to the mean LFQ intensity of all symmetric (for first three bars) or asymmetric (for last four bars) pulldown conditions analyzed. Error bars, 95% confidence interval of mean; diamonds, individual values of three independent nucleosome pulldown experiments, proteins color-coded as in panels (A) and (D). **(C)** Western Blot analysis of nucleosome pulldowns for select H3K4me3 and H3K27me3 binding proteins as well as HA-tagged KAT6B as bivalency-enriched protein. For the latter, NE of E14 ESCs stably expressing FLAG-HA-KAT6B from its endogenous locus was used for pulldown experiments. Data shown are representative of three independent pulldown experiments. **(D)** Same Volcano plot analysis as in (A), this time highlighting proteins specifically enriched on or depleted from bivalent nucleosomes. The C-terminal tags present on histone H3 in asymmetric nucleosomes did not significantly affect binding of proteins enriched on bivalent nucleosomes (see Figure S2B). Gold, KAT6B subunits; orange, SRCAP subunits; purple, other highlighted bivalent binders; teal, TAF subunits; blue, INO80 subunits; dark blue, zinc finger proteins; green and read, as in panel (A). **(E)** Model illustrating how bivalent nucleosomes support poising of genes by recruiting H3K27me3 binders as well as bivalency-specific binders, but not binders of the active H3K4me3 mark. See also Figure S3 and Table S3.

Strikingly, we further identified several proteins that were specifically enriched on asymmetric bivalent but not monovalent symmetric H3K4me3 or H3K27me3 nucleosomes (Figures 3B, 3D, S3A, Table S3), suggesting that asymmetric bivalent nucleosomes recruit specific factors through multivalent interactions involving both marks. These factors included two chromatin modifying complexes, the histone acetyltransferase KAT6B (also known as MORF) and the histone chaperone SRCAP. In addition to these complexes, we also found RNA binding proteins including ZRANB2, SRSF2, and PUM1, metabolic enzymes such as acetyl-CoA carboxylase, propionyl-CoA carboxylase, and 3-methylcrotonyl-CoA carboxylase, and transcription factors including MITF, TFE3, and TFEB to be specifically enriched on bivalent but not monovalent nucleosomes (Figures 3A–3C and S3A, Table S3). We also observed depletion of factors from bivalent nucleosomes, most prominently members of the INO80 complex, several zinc finger proteins, and general transcription factor subunits (Figure 3D and S3A, Table S3). However, the majority of these depleted proteins displayed favorable interactions with the asymmetric unmodified control nucleosomes (Figure S3B), potentially confounding accurate assessment of the role of the bivalent marks in recruitment of these factors.

Taken together, these findings suggest that the asymmetric bivalent histone mark combination directly promotes poising by recruiting repressive and bivalency-specific factors but excluding active factors, thereby maintaining a repressed but plastic state (illustrated in Figure 3E). To examine whether the differential recruitment of H3K4me3 and H3K27me3 binders was recapitulated *in vivo,* we analyzed the genome-wide distribution of TAF3 and CBX7 (Liu et al., 2011; Morey et al., 2013). CBX7 was found enriched at bivalent promoters within the Hoxd cluster and on a genome-wide scale in ESCs, whereas TAF3 was largely absent from bivalent promoters, despite presence of H3K4me3 (Figures 4A-C). Conversely, TAF3 was enriched at promoters monovalently marked with H3K4me3 (Figures 4A–C). These data are in agreement with the *in vitro* nucleosome pulldown experiments and support a histone mark-based poising mechanism via recruitment of specific repressive complexes and exclusion of active binders at bivalent domains.

**Figure 4.**
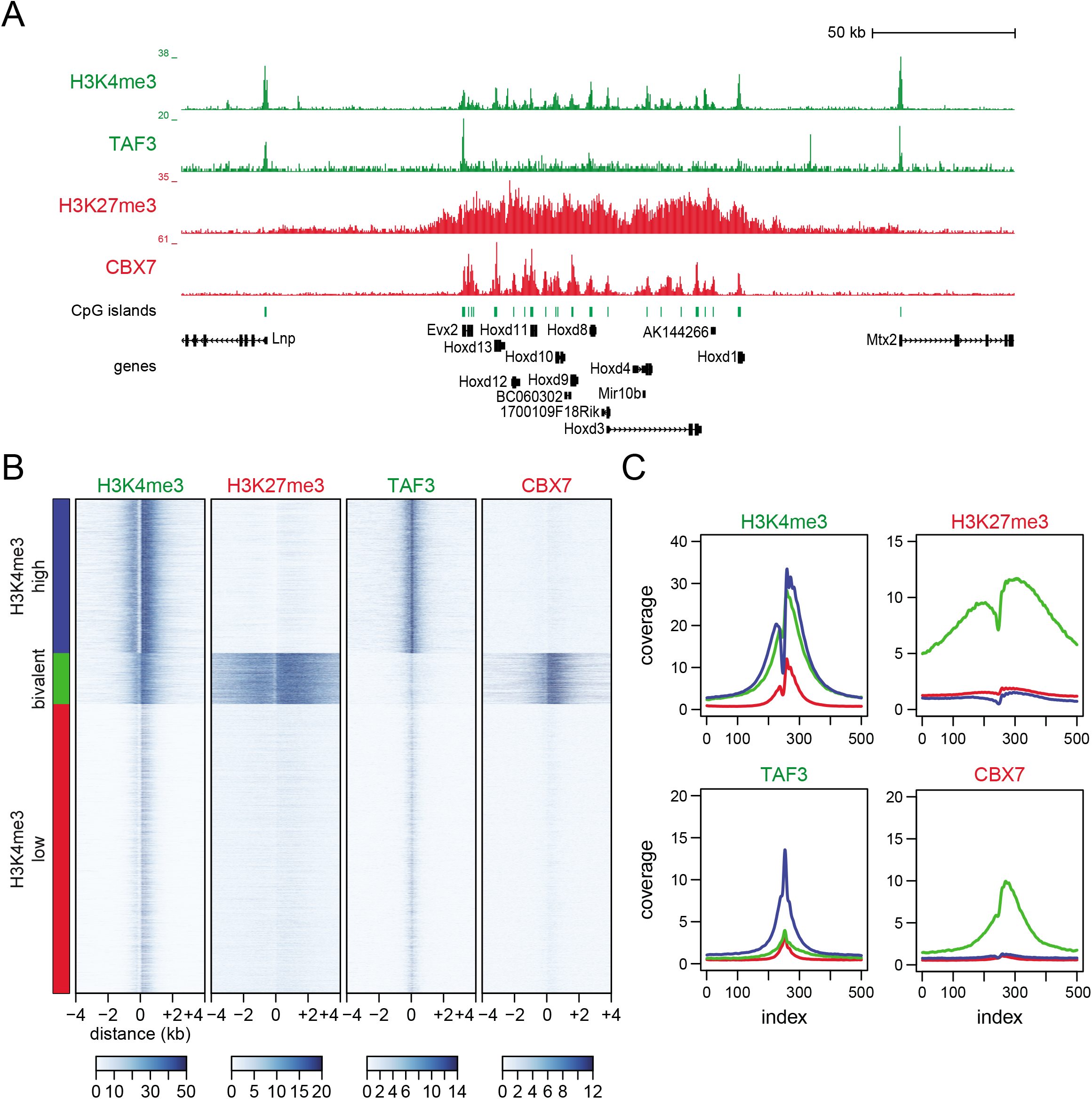
Bivalent nucleosomes in ESCs recruit CBX7 but not TAF3. **(A)** Genome browser snapshot showing distribution of H3K4me3, TAF3, H3K27me3, and CBX7 over the Hoxd cluster and adjacent active genes. For all panels, TAF3 ChIPseq data is from Liu et al. (2011) and data for CBX7 from Morey et al. (2013). Positions of CpG islands and genes are indicated below the tracks. **(B)** Heat maps of H3K4me3, H3K27me3, TAF3, and CBX7 ChIP-seq read densities centered around transcriptional start sites (TSS) and clustered as indicated, by high or low enrichment of H3K4me3 and H3K27me3. Note presence of TAF3 on H3K4me3-positive but not bivalent TSS, whereas CBX7 is bound to the latter. **(C)** Representation of data as average density profiles around TSS clustered into H3K4me3-positive (blue), bivalent (green), and H3K4me3-negative (red) TSS as in (B).

To understand how the bivalency-specific factors identified in our nucleosome pulldown assays (Figure 3D) contribute to the regulation of developmental genes at bivalent promoters *in vivo,* we focused on the chromatin modifying complexes SRCAP, a histone chaperone complex responsible for incorporating H2A.Z–H2B dimers into nucleosomes (Clapier et al., 2017; Giaimo et al., 2019; Mizuguchi et al., 2004), and KAT6B, a multi-subunit histone acetyltransferase that, along with its paralog KAT6A/ MOZ, has been implicated in acetylation of H3K9, H3K14, and H3K23 (Huang et al., 2016; Klein et al., 2019; Yang, 2015). Co-enrichment of several core subunits suggested recruitment of intact, functional complexes for both SRCAP (subunits DMAP1, YEATS4/GAS41, and VPS72) and KAT6B (subunits BRPF1/3, MEAF6, and ING5; Figures 3B, 3D, S3A). Whereas SRCAP has been previously linked to bivalency through presence of H2A.Z at bivalent domains in ESCs (Creyghton et al., 2008; Ku et al., 2012) and the requirement of H2A.Z for ESC differentiation (Creyghton et al., 2008; Hu et al., 2013b), the role of KAT6B and its acetylation marks at bivalent domains is unclear. We therefore asked whether KAT6B is recruited to bivalent domains in ESCs and whether KAT6B contributes to regulation of developmental genes during differentiation.

### Bivalency promotes recruitment of KAT6B to developmental genes in ESCs

To determine whether KAT6B is recruited to bivalent promoters in ESCs, we generated ESC lines expressing N-terminally FLAG- and HA-tagged KAT6B from the endogenous Kat6b gene via CRISPR/Cas9-based genome editing (Figures S4A and S4C) and performed chromatin immunoprecipitation (ChIP) using the HA epitope. In line with our nucleosome pulldown data (Figure 3D), KAT6B was bound to bivalent promoters, but not inactive promoters or gene bodies of bivalent genes (Figure 5A). In addition, KAT6B was also found at promoters of active genes (Figure 5A). To test whether recruitment of KAT6B to bivalent promoters in ESCs is dependent on the bivalent state, we introduced a point mutation that renders MLL2, the methyltransferase responsible for placing H3K4me3 at bivalent domains (Denissov et al., 2014; Hu et al., 2013a), catalytically inactive (Figures 5A, S4B, S4C). Concomitant with the reduction in H3K4me3, levels of KAT6B were markedly reduced at bivalent promoters in MLL2 Y2688A-expressing ESCs (Figure 5A). These findings indicate that recruitment of KAT6B to bivalent promoters is promoted by the bivalent modification state, in line with our nucleosome pulldown data.

**Figure 5.**
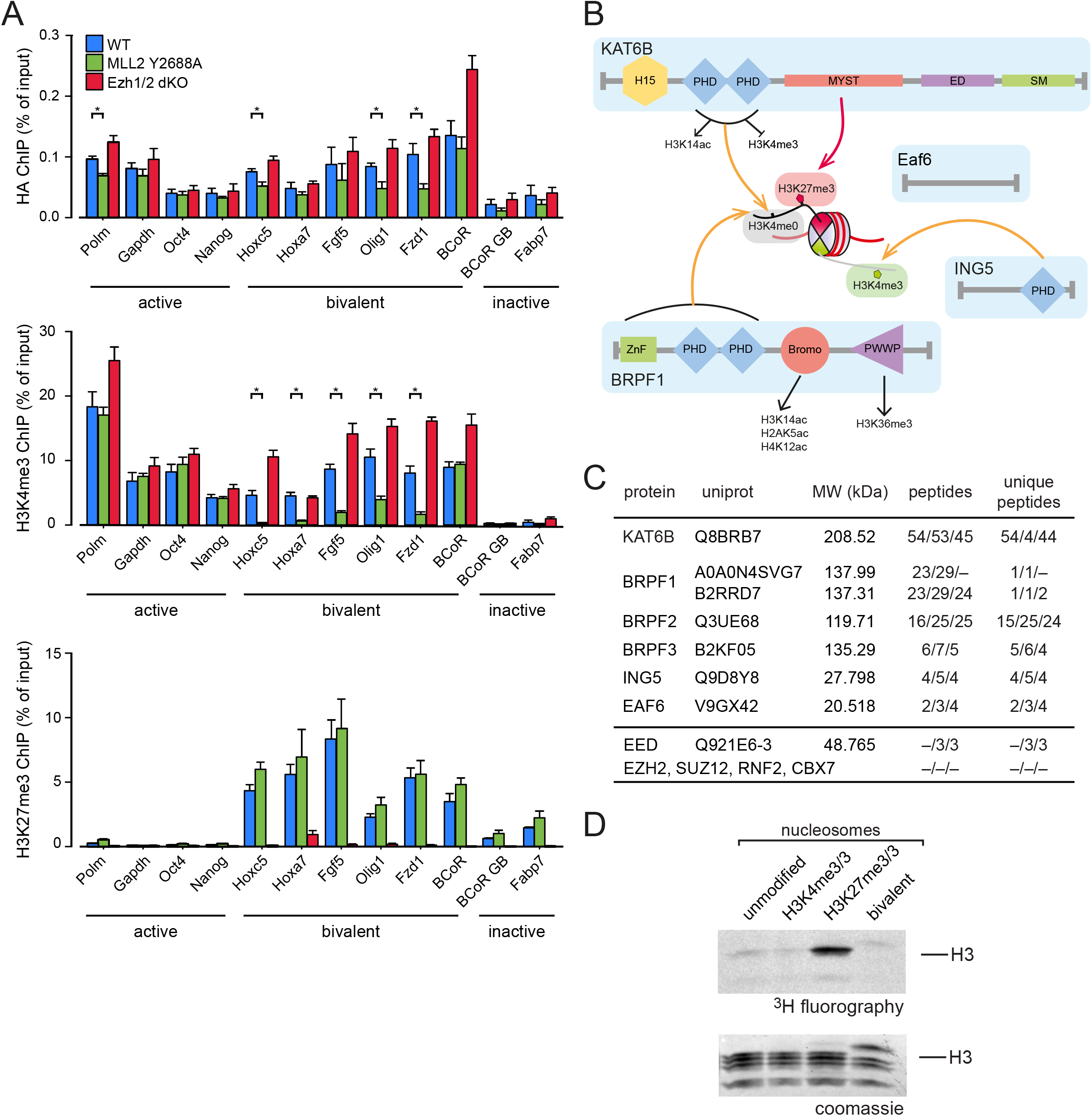
Bivalency promotes recruitment of KAT6B to developmental genes in ESCs. **(A)** ChIP-qPCR analysis with HA tag (upper panels), H3K4me3 (middle panels), and H3K27me3 (lower panels) antibodies for select active, bivalent, and inactive genes in E14 ESCs expressing FLAG-HA tagged KATB from its endogenous locus (blue bars), and in ESC lines with additional mutation in MLL2 (catalytically inactive MLL2 Y2688A, green bars) or knockout of EZH2 and EZH1 (red bars). Plotted are means and SEM of four independent ChIP experiments. Statistically significant differences (p ≤ 0.05, Student’s t test) in HA and H3K4me3 ChIP for control and MLL2 Y2688A background are highlighted by asterisks. **(B)** Domain composition of KAT6B complex and proposed model for its interaction with bivalent nucleosomes based on previously described histone mark interaction and data obtained here. **(C)** MS analysis of FLAG purification of FLAG-HA-KAT6B from NE of E14 ESC stably expressing tagged KAT6B from its endogenous locus. Note co-purification of all known KAT6B complex subunits but absence of PRC1 and PRC2 subunits. **(D)** Histone acetyltransferase (HAT) assay with KAT6B complex purified from ESC NE as in (C). KAT6B complex was incubated with the indicated nucleosome species and activity was monitored via ^3^H incorporation from radiolabeled acetyl-CoA. Equal loading was verified by Coomassie stain as shown below. Experiment shown representative of three independent HAT assays. See also Figure S4.

We next sought to clarify the mechanisms underlying recruitment of KAT6B to bivalent promoters. The KAT6B complex comprises a host of well-characterized histone mark binding domains (Figure 5B). The PHD finger of ING5 has been shown to bind H3K4me3 (Champagne et al., 2008), whereas the double PHD fingers of both KAT6B (Ali et al., 2012; Klein et al., 2017; Qiu et al., 2012) and BRPF1/2 (Lalonde et al., 2013; Qin et al., 2011) bind histone H3 tails with unmodified but not trimethylated H3K4. These interactions provide a mechanism for recognition of the asymmetric H3K4me3 feature contained in bivalent nucleosomes (illustrated in Figure 5B).

Given failure of asymmetric H3K4me3 nucleosomes to enrich KAT6B in pulldown assays *in vitro* (Figures 1C and 3B), recruitment of KAT6B requires H3K27me3 at least *in vitro* (Figures 3B–3D). However, KAT6B does not contain any known H3K27me3 binding domains. Enok, the *Drosophila* ortholog of KAT6B, has been shown to interact with PRC1 (Kang et al., 2017; Kang et al., 2015; Strübbe et al., 2011), which could mediate interaction with H3K27me3 in a KAT6B-PRC1 complex. However, we did not observe interaction of murine KAT6B with PRC1 (Figure 5C). Moreover, RING1B binding at bivalent promoters in ESCs was diminished upon EZH1/2 double knockout (Figure S4D), whereas KAT6B binding remained largely unaffected (Figure 5A), further arguing against an essential role of PRC1 in recruiting KAT6B in mouse ESCs.

Given that KAT6B acetylates H3K23 (Huang et al., 2016; Klein et al., 2019; Simó-Riudalbas et al., 2015), we reasoned that the catalytic MYST domain of KAT6B might interact with a H3K27me3 mark present adjacent to H3K23. Indeed, when performing histone acetyltransferase assays with KAT6B complex purified from ESCs, KAT6B was markedly more active towards H3K27me3 nucleosomes than towards unmodified or H3K4me3-modified nucleosomes (Figure 5D). These findings suggest that the catalytic MYST domain interacts with H3K27me3 as part of substrate recognition, providing a transient H3K27me3 binding site that promotes recruitment of KAT6B to bivalent nucleosomes (illustrated in Figure 5B). After acetylation, H3K23ac could interact with the double PHD finger of KAT6B or the bromodomain of BRPF1 (illustrated in Figure 5B), providing additional multivalent interaction sites for the KAT6B complex on H3K23ac-marked bivalent nucleosomes. To test the role of H3K27me3 in recruitment of KAT6B *in vivo,* we introduced an EZH1/2 double knockout into the ESC lines harboring tagged KAT6B (Figures S4B and S4C). Despite absence of H3K27me3, levels of KAT6B at bivalent promoters remained largely unchanged (Figure 5A). These observations suggest that additional interactions at bivalent domains *in vivo* could mask the dependence of recruitment on H3K27me3 that is clearly discernible in nucleosome pulldown assays *in vitro*. These interactions could involve other histone marks such as H3K14ac, which is present on both active and bivalent promoters in ESCs (Figure S5C).

### KAT6B is required for neuronal differentiation

Having established that KAT6B is recruited to bivalent domains *in vivo*, we next asked whether KAT6B regulates expression of bivalent genes in ESCs and during differentiation. To this end, we generated knockout ESC lines for KAT6B and its paralog KAT6A using CRISPR/Cas9 genome editing (Figures S5A and S5B). KAT6A/B double knockout ESC lines exhibited diminished H3K23ac, whereas single knockouts displayed only partial reductions in H3K23ac (Figures 6A and S5B). In contrast, acetylation of H3K14 remained largely unchanged (Figure S5C), indicating that KAT6A/B are essential for H3K23 but not H3K14 acetylation in mouse ESCs. H3K4me3 and H3K27me3 were unaffected by knockout of KAT6A/B (Figure S5C). Knockout of KAT6A/B did not affect proliferation or maintenance of ESC identity (Figure 6B). Moreover, despite diminished H3K23ac at their promoters, expression of housekeeping genes was unaffected in KAT6A/B knockout ESCs (Figure S5D), suggesting that KAT6A/B and the H3K23ac mark are dispensable for expression of active genes in ESCs. Furthermore, we did not observe derepression of bivalent genes in KAT6A/B knockout ESCs (Figure 6D), indicating that KAT6B is not required for maintaining their repression in ESCs.

**Figure 6.**
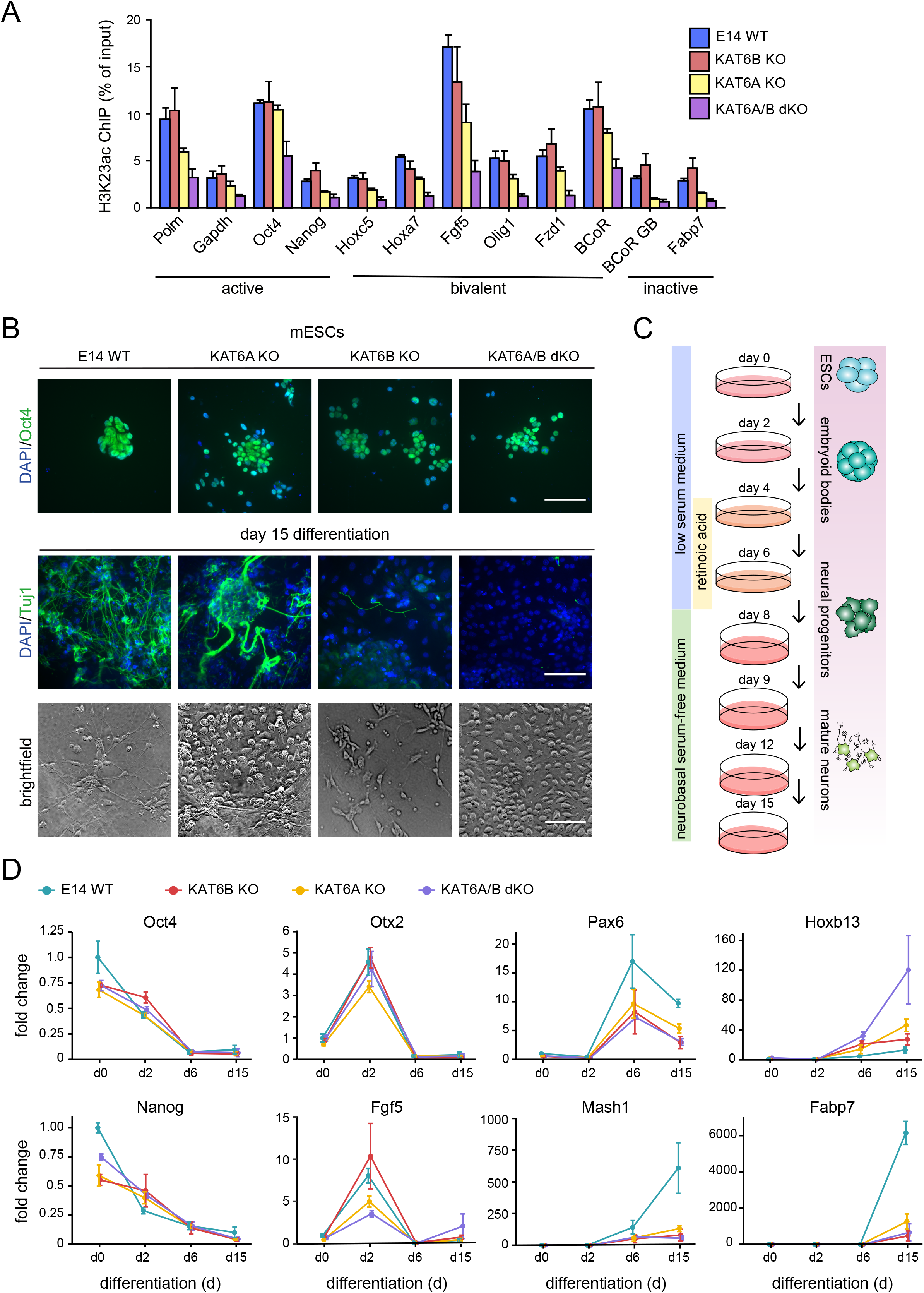
KAT6B is required for neuronal differentiation. **(A)** ChIP-qPCR analysis with H3K23ac antibody for select active, bivalent, and inactive genes in E14 ESCs (blue bars), KAT6B knockout (red bars), KAT6A knockout (yellow bars), and KAT6A/B double knockout (purple bars) E14 ESCs. Plotted are means and SEM of four independent ChIP experiments. **(B)** Oct4 and Tuj1 immunofluorescence staining of ESCs before and after the neuronal differentiation protocol outlined in (C). Note that neuronal protrusions positive for Tuj1 are formed in differentiated E14 control ESCs but not in KAT6 knockout ESCs. Brightfield images further show presence or absence of protrusions. Images shown are representative of three independent differentiation experiments. Scale bars, 100 μm. **(C)** Outline of neuronal differentiation protocol. **(D)** RT-qPCR analysis of mRNA expression changes in select pluripotency, early differentiation, and neuronal marker genes in E14 ESCs and KAT6 knockout ESC lines, expressed as fold changes relative to day 0 and normalized to expression in E14 ESC controls on day 0. Data are represented as mean and SEM of three independent differentiation experiments. See also Figure S5.

To test whether KAT6B is instead required to regulate proper induction or silencing of bivalent genes upon differentiation, we carried out neuronal differentiation towards the glutamatergic neuronal lineage via formation of embryoid bodies (EBs) and neural progenitor cells (Bibel et al., 2007) (Figure 6B–6D). After 15 days, formation of mature neurons was clearly evident for wild-type E14 ESCs by the presence of networks of neuronal protrusions that were positive for the neuron-specific class III ß-tubulin Tuj1 (Figure 6B). Strikingly, KAT6A/B single and double knockouts failed to generate mature neurons (Figure 6B), indicating diminished differentiation potential of KAT6A/B knockout ESCs. To determine how loss of KAT6A/B compromises differentiation, we analyzed expression of pluripotency and differentiation marker genes. Whereas downregulation of pluripotency factors Oct4 and Nanog was largely unaffected in KAT6A/B knockout cells, expression of early differentiation markers Otx2 and Fgf5 was misregulated after 2 days of differentiation (Figure 6D). These genes feature bivalent promoters, indicating a role for KAT6B and KAT6A in regulation of bivalent genes at the onset of differentiation. Moreover, KAT6A/B knockout lines failed to upregulate early and late neuronal markers such as Pax6, Mash1, and Fabp7 (Figure 6D), further supporting a pivotal role of KAT6B and KAT6A in the regulation of bivalent developmental genes during differentiation. Interestingly, Hoxb13 was excessively upregulated in KAT6A/B knockout cell lines (Figure 6D), indicating a failure to also properly curtail expression of some bivalent genes in addition to promoting expression of others. Taken together, these findings indicate that KAT6B is required for proper differentiation of ESCs through regulation of bivalent developmental genes.

## Discussion

Bivalent domains have been posited to poise developmental genes for timely expression or terminal repression upon differentiation while keeping their expression repressed in ESCs (Azuara et al., 2006; Bernstein et al., 2006; Mikkelsen et al., 2007; Voigt et al., 2013). However, it has remained unclear whether and how the histone modifications present at bivalent domains directly cause—or contribute to— establishment of a poised state. Our findings reveal a histone mark-based poising mechanism at bivalent domains and highlight how nucleosomal asymmetry regulates histone mark readout and function through allovalency, uncovering a novel regulatory mechanism in the recruitment of chromatin binders. We show that the bivalent histone marks themselves are sufficient to set up a poised state, revealing a direct role for the bivalent modification state in regulating expression of developmental genes. We propose that bivalent nucleosomes promote poising by recruiting the repressive H3K27me3 binders PRC1 and PRC2 along with bivalency-specific binders, while excluding active mark binders such as TAF3, preventing gene activation despite presence of H3K4me3 (Figure 3E).

The asymmetric modification state of bivalent nucleosomes appears to be crucial for the differential recruitment of active and repressive binders at bivalent domains. For symmetrically modified nucleosomes, allovalent effects associated with the presence of two equivalent binding sites in close proximity boost interaction with mark binding proteins (see Figure 2B). These effects support efficient recruitment of even lowly abundant binding proteins at symmetric nucleosomes, whereas recruitment of highly abundant binders would be favored at asymmetric nucleosomes, as their abundance overcomes the reduced affinity associated with asymmetric modification (see Figure 2C). Accordingly, in nucleosome pulldown assays recruitment of H3K4me3 binders such as TAF3 was robust for symmetric but not asymmetric nucleosomes, whereas the more highly abundant H3K27me3 binders were efficiently recruited also at asymmetric nucleosomes (Figure 2A and 2D). Consistent with these findings and our proposed model, bivalent promoters in ESCs are bound by PRC complex members such as CBX7 but fail to recruit TFIID, as shown for TAF3 (Figure 4A–C; Liu et al., 2011) and TBP (Ku et al., 2012).

Beyond bivalent domains, nucleosomal asymmetry of H3K4me3 could control global transcriptional output by regulating recruitment of TFIID and potentially of other H3K4me3 binders that facilitate transcription. Strikingly, H3K4me3 abundance at promoters in mouse ESCs, when calibrated and normalized to biologically meaningful modification densities using internal standards in ChIP-seq, correlates with RNA expression in a sigmoidal fashion (Grzybowski et al., 2015). Promoters with H3K4me3 densities corresponding to asymmetric modification exhibit low expression, whereas expression is markedly increased above the asymmetric-symmetric transition (Grzybowski et al., 2015), potentially reflecting more efficient recruitment of H3K4me3 binders to symmetrically modified nucleosomes due to allovalent effects. Interestingly, based on recent cryo-EM structures of human TFIID (Patel et al., 2018), it has been proposed that recognition of promoter nucleosomes by TFIID could precede engagement of promoter DNA by TBP (Bhuiyan and Timmers, 2019; Patel et al., 2018), supporting a crucial role for the TAF3-H3K4me3 interaction in TFIID recruitment. Similar to multivalency, a central principle governing affinity and specificity of chromatin-based interactions via combinatorial readout of histone marks, DNA, and RNA (see e.g. Patel and Wang, 2013; Ruthenburg et al., 2007; Su and Denu, 2016), allovalency could therefore be crucial to regulating the recruitment and functional output of chromatin complexes, especially for lowly abundant, narrowly distributed marks such as H3K4me3 and their cognate, lowly abundant binders.

In addition to curbing recruitment of H3K4me3 binders, asymmetric bivalent nucleosomes enrich PRC1 and PRC2 complexes. Given the central role of binding proteins in mediating histone mark function, the proteins recruited to bivalent nucleosomes likely set up or promote a poised state. Several mechanisms have been proposed for PRC-mediated repression, including compacting chromatin, curbing productive elongation by RNAPII, promoting a poised, exclusively S5-phosphorylated state of RNAPII, and blocking acetylation of H3K27 through PRC2-mediated H3K27 methylation (Aranda et al., 2015; Schuettengruber et al., 2017; Simon and Kingston, 2013; Voigt et al., 2013). Given that allovalent effects at symmetric H3K27me3 nucleosomes enhance recruitment especially of PRC1 (Figures 1D, 3A, 3B, S3A), it is conceivable that these modes of repression are attenuated at asymmetric bivalent nucleosomes, establishing a poised rather than fully repressed state. Indeed, removal of asymmetric H3K4me3 at bivalent domains via knockout of MLL2 increases levels of H3K27me3, likely converting asymmetric bivalent to symmetric H3K27me3 nucleosomes, leading to increased recruitment of PRC2 and PRC1 and, in turn, reduced promoter accessibility and levels of Ser5-phosphorylated RNAPII at bivalent domains (Mas et al., 2018). These findings provide additional support for a key role of nucleosomal asymmetry in setting up a poised rather than fully repressed state at bivalent domains.

Interestingly, asymmetric bivalent nucleosomes displayed preference for PRC2 complexes containing EPOP and EZHIP (Figures 3A and S3A). EPOP and EZHIP have been shown to reduce H3K27me3 levels by competing with binding of PRC2 auxiliary subunits such as JARID2 that confer more potent repressive activity to PRC2 (Beringer et al., 2016; Liefke et al., 2016; Ragazzini et al., 2019) and, in the case of EZHIP, also by directly inhibiting PRC2 activity (Hübner et al., 2019; Jain et al., 2019). The bivalency-specific recruitment of EPOP and EZHIP may therefore further contribute to poising by curbing PRC2 activity. In agreement with our nucleosome pulldown data, EPOP has been shown to localize to bivalent domains in ESCs (Beringer et al., 2016; Liefke et al., 2016; Liefke and Shi, 2015).

In addition, we show that bivalent nucleosomes recruit factors that specifically interact with bivalent nucleosomes (Figures 3B–D, S3A), providing a set of candidates for novel mediators of poising, functioning alongside PRC1 and PRC2 at bivalent domains (Figure 3E). In contrast to the plant proteins EBS and SHL, which were recently shown to recognize both H3K27me3 and H3K4me3 in a mutually exclusive fashion (Qian et al., 2018; Yang et al., 2018), the bivalency-specific binders identified here simultaneously and multivalently engage both marks to achieve recruitment to bivalent domains. Interestingly, besides KAT6B and SRCAP, most identified factors lack known histone binding domains, suggesting involvement of novel binding domains or recognition of composite binding surfaces generated by binding of PRCs, SRCAP, or KAT6B to bivalent nucleosomes.

To clarify how the novel bivalency-specific factors contribute to the regulation of developmental genes at bivalent promoters we focused our analysis on KAT6B. We show that the complex is recruited to bivalent domains in ESCs (Figure 5A) and crucial to regulating expression of bivalent domains during differentiation (Figure 6). The KAT6B complex features well-characterized histone mark binding domains, suggesting a model for the recruitment of KAT6B to bivalent nucleosomes (Figure 5B). Multivalent interactions involving additional binding domains present in the complex could further contribute to its recruitment to bivalent domains as well as active promoters in ESCs *in vivo*. Our findings reveal that KAT6B contributes to maintaining a poised chromatin state while being dispensable for active transcription, potentially by maintaining accessibility of bivalent promoters through placement of H3K23ac or other means. In support of such a role in chromatin accessibility, KAT6B has recently been shown to facilitate interaction of OCT4 and NANOG with chromatin in ESCs (Cosentino et al., 2019). Moreover, KAT6B-mediated H3K23ac could further contribute to poising by modulating interaction of PRC1 and PRC2 with H3K27me3, affecting recruitment of repressive complexes.

This study provides insight into the role of nucleosomal asymmetry and bivalency-specific multivalency in regulating recruitment of binding proteins to bivalent domains. Although the exact roles of Polycomb complexes and other factors recruited to bivalent domains remain to be uncovered, our data provides evidence that the bivalent histone marks themselves are sufficient to set up a poised state at bivalent domains. The recruitment of KAT6B and SRCAP to bivalent nucleosomes highlights the potential significance of additional layers of histone-based regulation at bivalent nucleosomes. Adding to the roles exerted by the ‘core’ bivalent modification comprised of H3K4me3 and H3K27me3, CpG island promoters of developmental genes in ESCs appear to adopt a state of ‘extended bivalency’ featuring H3K4me3, H3K27me3, H2A ubiquitinylation, H2A.Z, H3.3, H3K14ac, H3K23ac, and possibly other marks. Unraveling how these different modules interact at bivalent domains to recruit chromatin complexes, promote poising, and regulate timely access to DNA will be pivotal to further refining our understanding of the mechanisms that control expression of developmental genes in ESCs.

## Acknowledgments

We are grateful to Till Bartke for helpful discussions and advice on nucleosome pulldown assays. We thank Tania Auchynnikava for help with initial MS analysis and Giulia Bartolomucci for technical support in generating knockout and knockin ESC lines. We are grateful to Dónal O’Carroll for critical reading of the manuscript. We thank Adrian Bird, Atlanta Cook, and Ken Sawin for fruitful discussions and insightful comments on this work. We also thank members of our labs for helpful discussions. This work was supported by the Wellcome Trust ([104175/Z/14/Z], Sir Henry Dale Fellowship to P.V., [091020] Equipment grant to J.R., and Multi-User Equipment grant [108504]) and through funding from the European Research Council (ERC) under the European Union’s Horizon 2020 research and innovation programme (ERC-STG grant agreement No. 639253 to P.V.). The Wellcome Centre for Cell Biology is supported by core funding from the Wellcome Trust [203149]. We are grateful to the Edinburgh Protein Production Facility (EPPF) for their support. The EPPF was supported by the Wellcome Trust through a Multi-User Equipment grant [101527/Z/13/Z]. Research in the Baubec Lab is supported by the Swiss National Science Foundation (#157488 and #180345) and the Swiss initiative in Systems Biology (SystemsX.ch).

## Author contributions

E.B. conceived, designed, performed, and analyzed experiments. M.W., K.W., K.M., V.M. and C.A. designed and performed experiments. K.M. and E.B. further analyzed and visualized data and generated figures. C.S. performed MS experiments and analysis. T.B. provided conceptual input, analyzed and supervised experiments, performed bioinformatic analysis, and secured funding. J.R. provided conceptual advice, analysis tools, and supervision. J.R. further secured funding. P.V. conceived the study, designed, performed, and analyzed experiments, wrote the manuscript with input from E.B. and T.B., supervised the project, and secured funding. All authors contributed to editing and revision of the manuscript.

## Supplemental Figure Legends

**Figure S1, related to Figure 1 and Figure 3.**
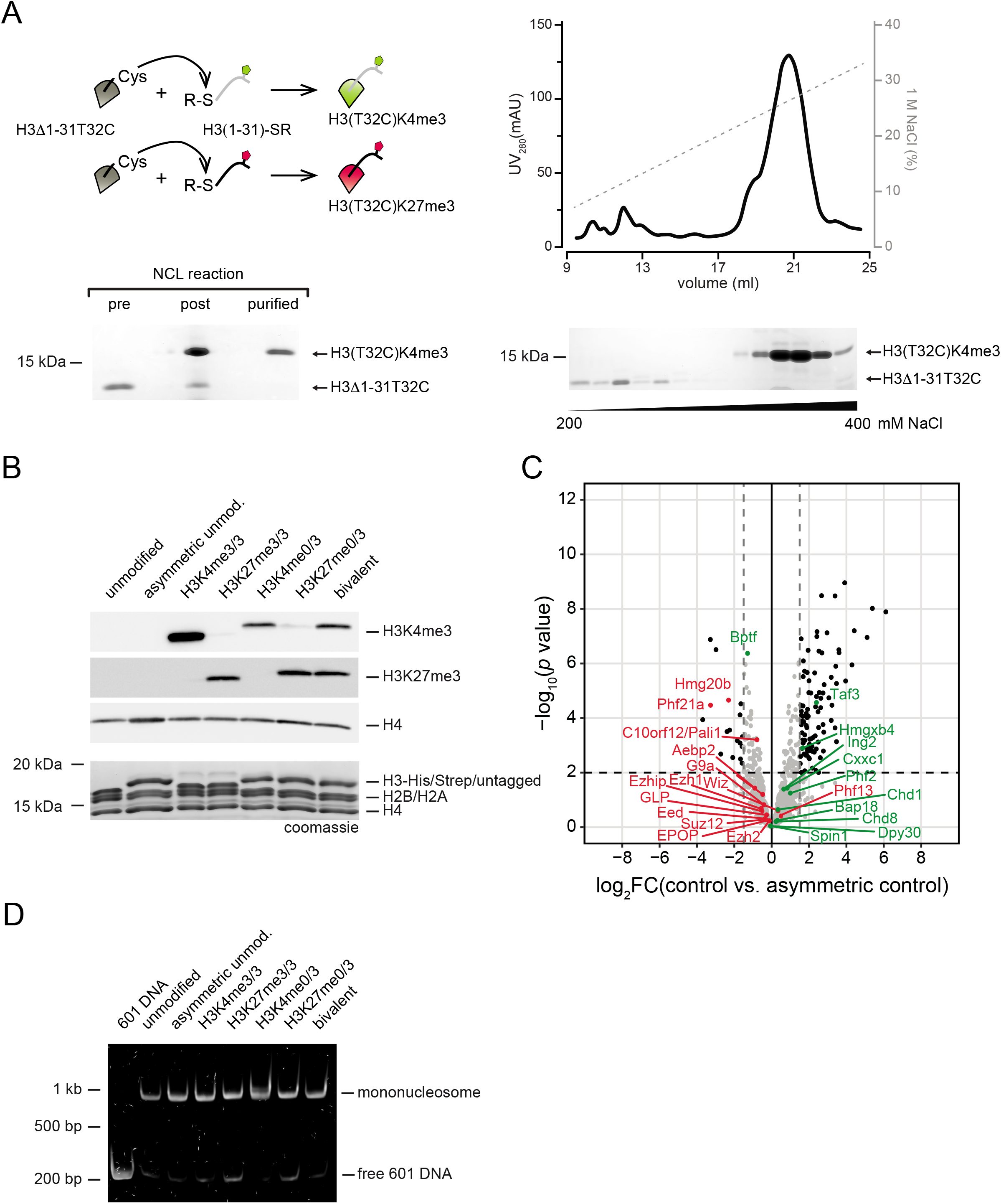
Generation of modified nucleosomes for pulldown experiments and control pulldowns. **(A)** Native chemical ligation (NCL) was used to generate modified histone H3 with either H3K4me3 or H3K27me3. *Top left panel,* schematic of NCL reaction involving truncated histone H3 lacking the first 31 amino acids after the initiator methionine (H3Δ1–31T32C) and a synthetic peptide spanning residues 1–31 of histone H3 and containing trimethylated lysine at the desired residue as well as a C-terminal thioester that is required for NCL. The reaction yields a full-length histone with the desired modification and a threonine-to-cysteine point mutation at position 32. For some experiments involving H3K27me3, T45 was used as the ligation site, resulting in a T45C point mutation in the final modified histone. *Bottom left panel,* Representative Coomassie gel analysis of NCL reaction for H3K4me3, showing unligated histone (pre), crude reaction mixture (post), and final ligated histone purified by cation exchange chromatography on a monoS column (purified). *Right panel,* representative cation exchange purification performed after NCL reactions to separate ligated from residual unligated histone. The elution profile of a linear NaCl gradient plotted as UV280 signal is shown along with a Coomassie stained SDS PAGE gel of fractions within the indicated range of the gradient. Peak fractions containing ligated histones were pooled and used for histone octamer assembly. **(B)** Analysis of symmetric and asymmetric octamers used for assembly of nucleosomes. Presence of H3K4me3 and H3K27me3 was verified by Western blotting with H3K4me3 and H3K27me3 antibodies. H4 was probed for as a loading control. Coomassie stain shows presence of histones H2B, H2A, and H4 along with untagged (symmetric octamers) and C-terminally Strep-or His-tagged histone H3 (asymmetric octamers). **(C)** Volcano plot analysis of LFQ MS data for pulldown experiments with unmodified nucleosomes containing C-terminally tagged His- and Strep-tagged histone H3 (used as controls for asymmetrically modified nucleosomes) compared to unmodified and untagged nucleosomes (used as controls for symmetrically modified nucleosomes) to assess the impact of the tags on binding to asymmetric nucleosomes. Means and *p* values are derived from three independent pulldown experiments. Black dots, significantly enriched or depleted proteins (log2 fold change ≤ 1.5 or ≥ 1.5, respectively, and *p* value ≤ 0.01). Known H3K4me3 and H3K27me3 binders are highlighted as in Figure 1. Note that presence of the C-terminal tags on histone H3 does not adversely affect binding of those factors. **(D)** Electrophoretic mobility shift assay (EMSA) performed to verify assembly of recombinant nucleosomes used for pulldown experiments shown in Figures 1 and 3. Mononucleosomes (nuc) were separated from free 601 DNA (601) on native 6% polyacrylamide gels in TGE buffer and visualized with SYBR safe stain.

**Figure S2, related to Figure 2.**
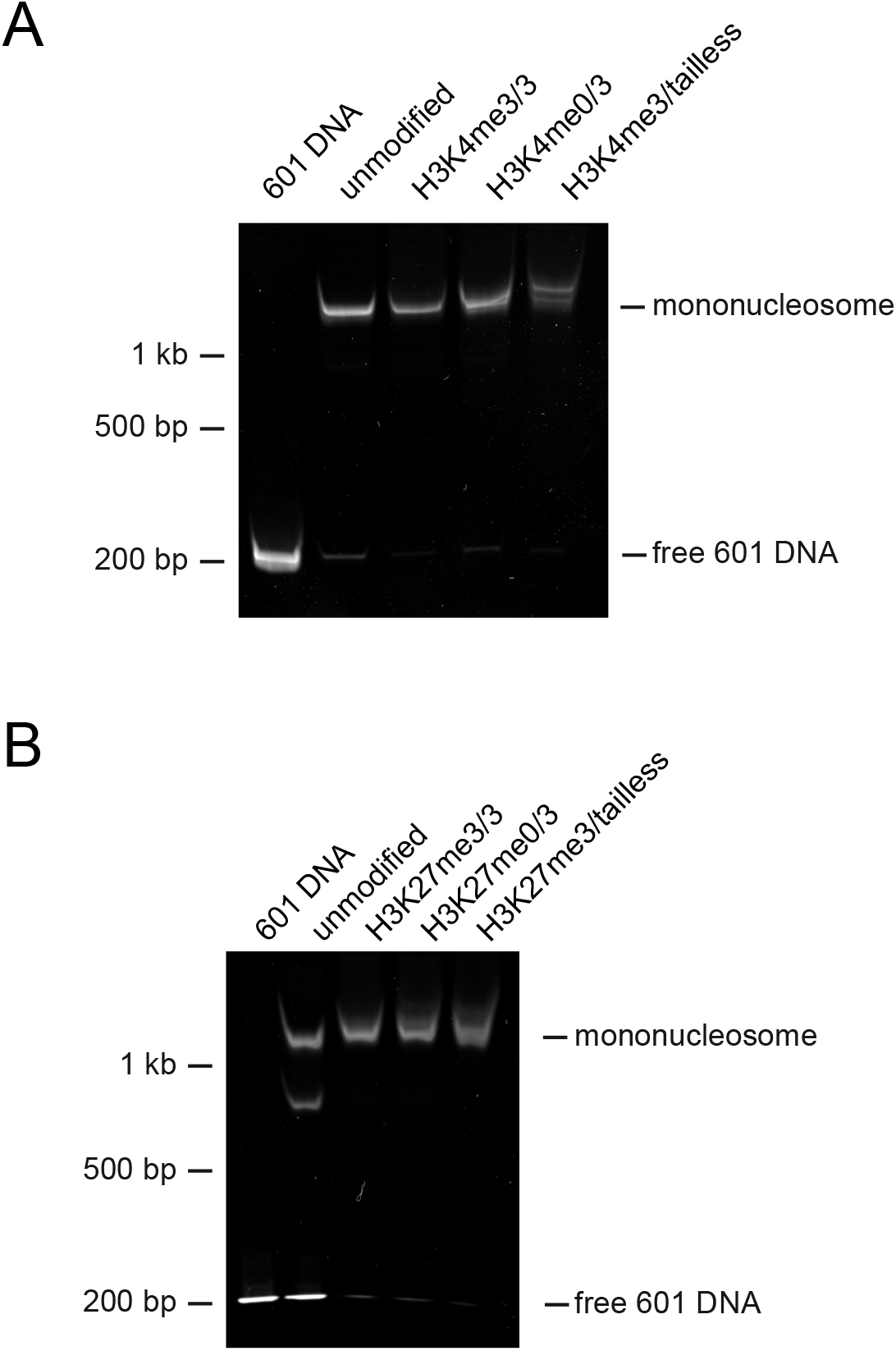
Nucleosome assembly for titration pulldowns. **(A,B)** EMSAs verifying assembly of recombinant nucleosomes used for pulldown experiments shown in Figure 2. Nucleosome preparations with H3K4me3 (A) and H3K27me3 (B) modifications were run on native 6% polyacrylamide gels in TGE buffer and visualized with SYBR safe stain.

**Figure S3, related to Figure 3.**
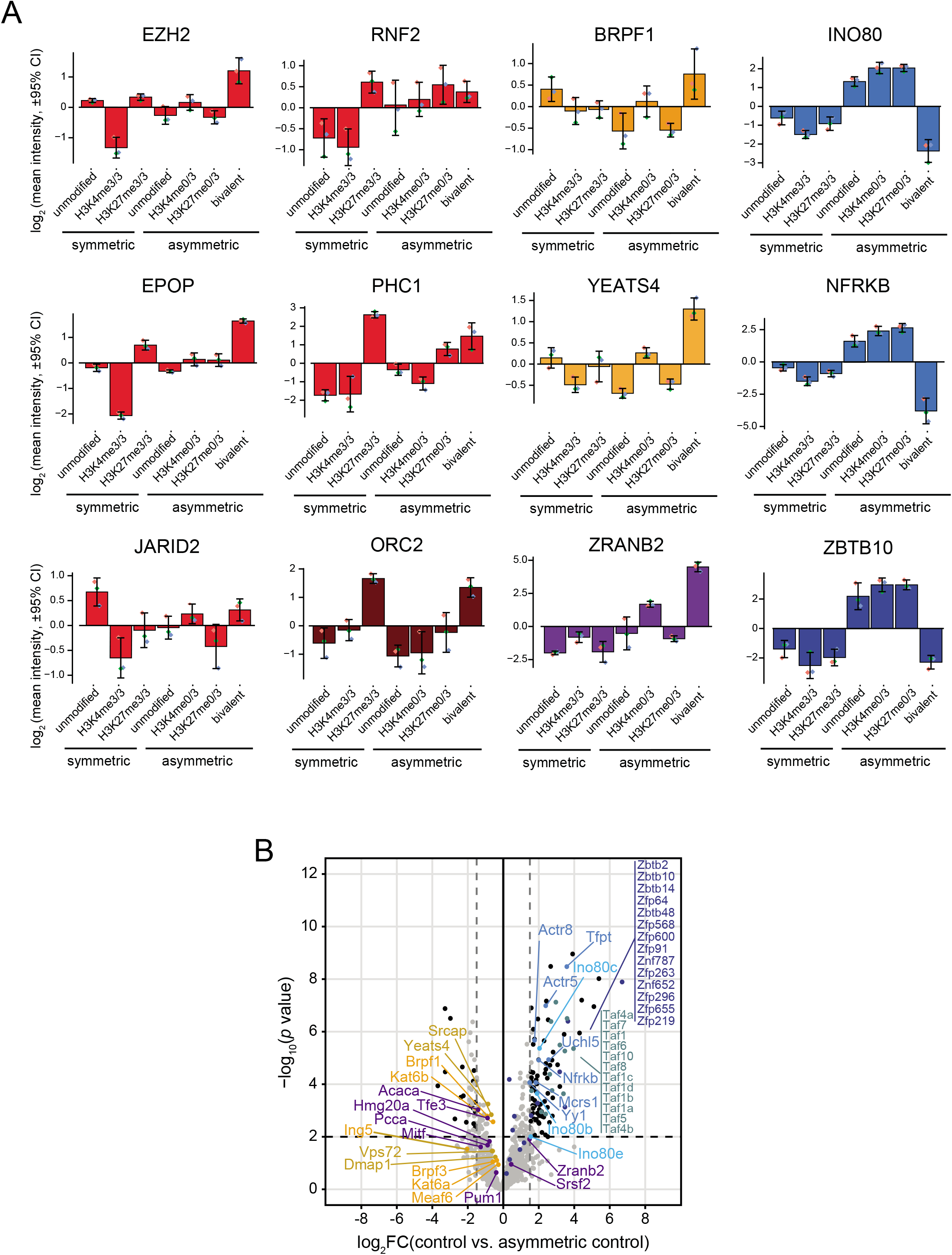
Nucleosome pulldown data for additional bivalent binders and control pulldowns. **(A)** Bars plots showing representative proteins specifically enriched or depleted on bivalent asymmetric nucleosomes. Plotted are fold changes of LFQ intensities relative to the mean LFQ intensity of all pulldown conditions analyzed. Error bars, 95% confidence interval of mean; diamonds, individual values of three independent nucleosome pulldown experiments, proteins color-coded as in Figure 3. **(B)** Volcano plot analysis of LFQ MS data for pulldown experiments with unmodified nucleosomes containing C-terminally tagged His- and Strep-tagged histone H3 (used as controls for asymmetrically modified nucleosomes) compared to unmodified and untagged nucleosomes (used as controls for symmetrically modified nucleosomes) to assess the impact of the tags on binding to asymmetric nucleosomes (same data as in Figure S1C). Proteins enriched or depleted on bivalent nucleosomes are highlighted as in Figure 3, showing that presence of the tags does not adversely affect binding of factors enriched on asymmetric bivalent nucleosomes, whereas several factors depleted from asymmetric bivalent nucleosomes exhibit favorable interactions with the tags.

**Figure S4, related to Figure 5.**
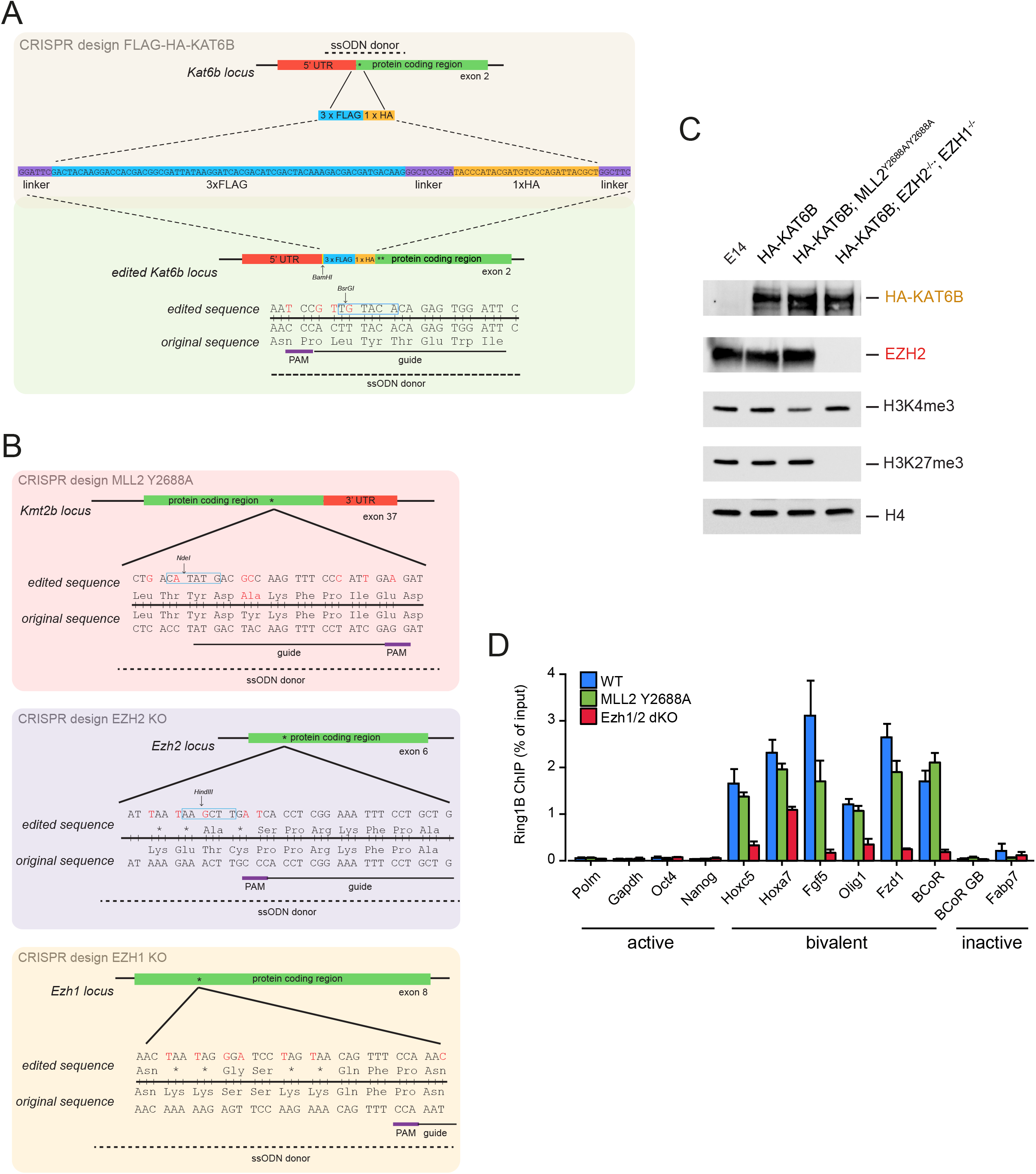
Generation and characterization of knockin and knockout cell lines and additional ChIP data. **(A)** Generation of E14 ESC lines expressing FLAG-HA-tagged KAT6B via CRISPR genome editing and homologous repair. **(B)** After successful generation of the ESC line expressing tagged KAT6B described in (A), additional CRISPR genome editing was performed to introduce a point mutation in the Kmt2b gene encoding the catalytically inactive Y2688A point mutation in MLL2 (top panel) as well as to generate a double knockout of EZH2 and EZH1 by introducing a series of premature stop codons in Ezh2 and Ezh1 genes (middle and bottom panels). **(C)** Western blot analysis on whole cell extracts of ESC lines generated, verifying presence of the HA tag on KAT6B, reduced levels of H3K4me3 in MLL2 Y2688A ESCs, and absence of EZH2 and H3K27me3 in EZH2/EZH1 double knockout ESCs. H4 was used as a loading control. Data shown are representative of two Western blot experiments. **(D)** ChIP-qPCR analysis with RING1B antibody for select active, bivalent, and inactive genes in E14 ESCs expressing FLAG-HA tagged KATB (blue bars), and cell lines with additional mutation in MLL2 (catalytically inactive MLL2 Y2688A, green bars) or knockout of EZH2 and EZH1 (red bars). Plotted are means and SEM of four independent ChIP experiments. Note that RING1B enrichment is diminished in absence of H3K27me3.

**Figure S5, related to Figure 6.**
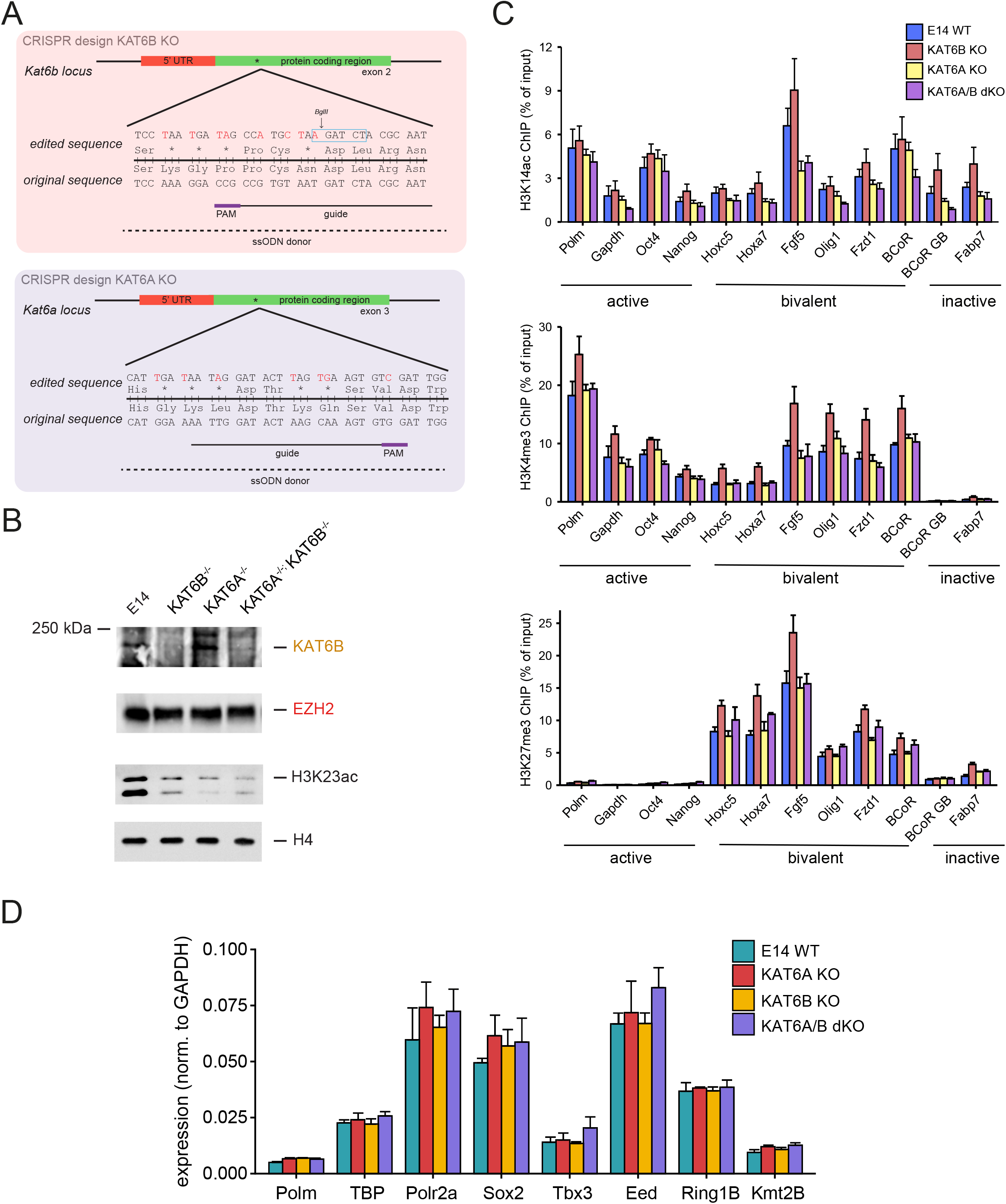
Generation and characterization of KAT6A/B knockout cell lines and additional ChIP and RT-qPCR data. **(A)** Generation of KAT6B and KAT6A knockout E14 ESCs via CRISPR genome editing and homologous repair to introduce a series of premature stop codons in Kat6b and Kat6a genes. **(B)** Western blot analysis confirming knockout of KAT6B and concomitant loss of H3K23ac in KAT6B and KAT6A knockouts, highlighting partial redundancy between both acetyltransferases. H4 and EZH2 were used as loading controls. Blots shown are representative of two independent experiments. **(C)** Additional ChIP-qPCR analysis of KAT6A/B knockout ESC lines with H3K14ac, H3K4me3, and H3K27me3 antibodies showing that these marks remain largely unchanged in the knockout ESC lines. Select active, bivalent, and inactive genes were analyzed in E14 ESCs (blue bars), KAT6B (red bars), KAT6A (yellow bars), and KAT6A/B double (purple bars) knockout ESCs. Plotted are means and SEM of four independent ChIP experiments. **(D)** RT-qPCR analysis of housekeeping gene expression in E14 ESCs and KAT6 knockout ESC lines, showing that housekeeping gene expression is not affected in knockout cell lines despite diminished H3K23ac levels in those cell lines. Expression was normalized to expression of Gapdh. Data are represented as mean and SEM of three independent experiments.

## Supplemental Tables

**Table S1.** Proteins significantly enriched or depleted for symmetric and asymmetric H3K4me3 nucleosomes.

| Complex | Direct binder | Significantly Enriched or Depleted Subunits |
| --- | --- | --- |
| Enriched for symmetric H3K4me3 nucleosomes: |  |  |
| SPIN1 | SPIN1 | SPIN1, SPINDOC |
| NURF | BPTF | BPTF, HMGXB4, BAP18, BAP18.1 |
| SIN3A | ING2 | ING2, BRMS1L, ARID4B, SUDS3, BRMS1, SAP30, SAP130, ARID4A, SAP30L, SIN3A, FAM60A |
| SET1 | CXXC1 | CXXC1, RBBP5, DPY30, SETD1A |
| TFIID | TAF3 | TAF3 |
| EMSY | JARID1A | SIN3B, JARID1A, GATAD1, EMSY, PHF12 |
| not assigned to complexes |  | ZGLP1, PHF2, MORC3, PHF13 (SPOC1), CHD8, PHF23, CHD6, DIDO1, JARID1B, CHD1 |
| Depleted for symmetric H3K4me3 nucleosomes: |  |  |
| PRC2 |  | EPOP, C10orf12/PALI1, EZH1, AEBP2, EED.1, SUZ12, EZH2, MTF2 |
| PRC1 |  | PCGF6 |
| G9A/GLP |  | G9A, GLP, WIZ |
| not assigned to complexes |  | PHF21A, HMG20B, BEND3, UHRF1, TAF1C, THAP1, BAZ2A, PRDM10, ZFP219, PAX6, BAZ2B, TOP3A, HIF1A |
| Enriched for asymmetric H3K4me3 nucleosomes*: |  |  |
| not assigned to complexes |  | ZRANB2, DNAJC9, MBD1 |

\*: not significant by adjusted p value.

**Table S2.** Proteins significantly enriched or depleted for symmetric and asymmetric H3K27me3 nucleosomes.

| Complex | Direct binder | Significantly Enriched or Depleted Subunits |
| --- | --- | --- |
| Enriched for symmetric H3K27me3 nucleosomes: |  |  |
| PRC1 | CBX7 | CBX7, PHC1 |
| ORC |  | ORC3, LRWD1, ORC2, ORC1 |
| not assigned to complexes |  | ARID4B, PHF2 |
| Depleted for symmetric H3K27me3 nucleosomes: |  |  |
| PRC2 |  | AEBP2 |
| not assigned to complexes |  | G2E3,ZBTB2. |
| Enriched for asymmetric H3K27me3 nucleosomes*: |  |  |
| not assigned to complexes |  | NFYB, NFYC, SIX4 |
| Depleted for asymmetric H3K27me3 nucleosomes: |  |  |
| not assigned to complexes |  | SURF6*, GNL3 |

\*: not significant by adjusted p value.

**Table S3.**
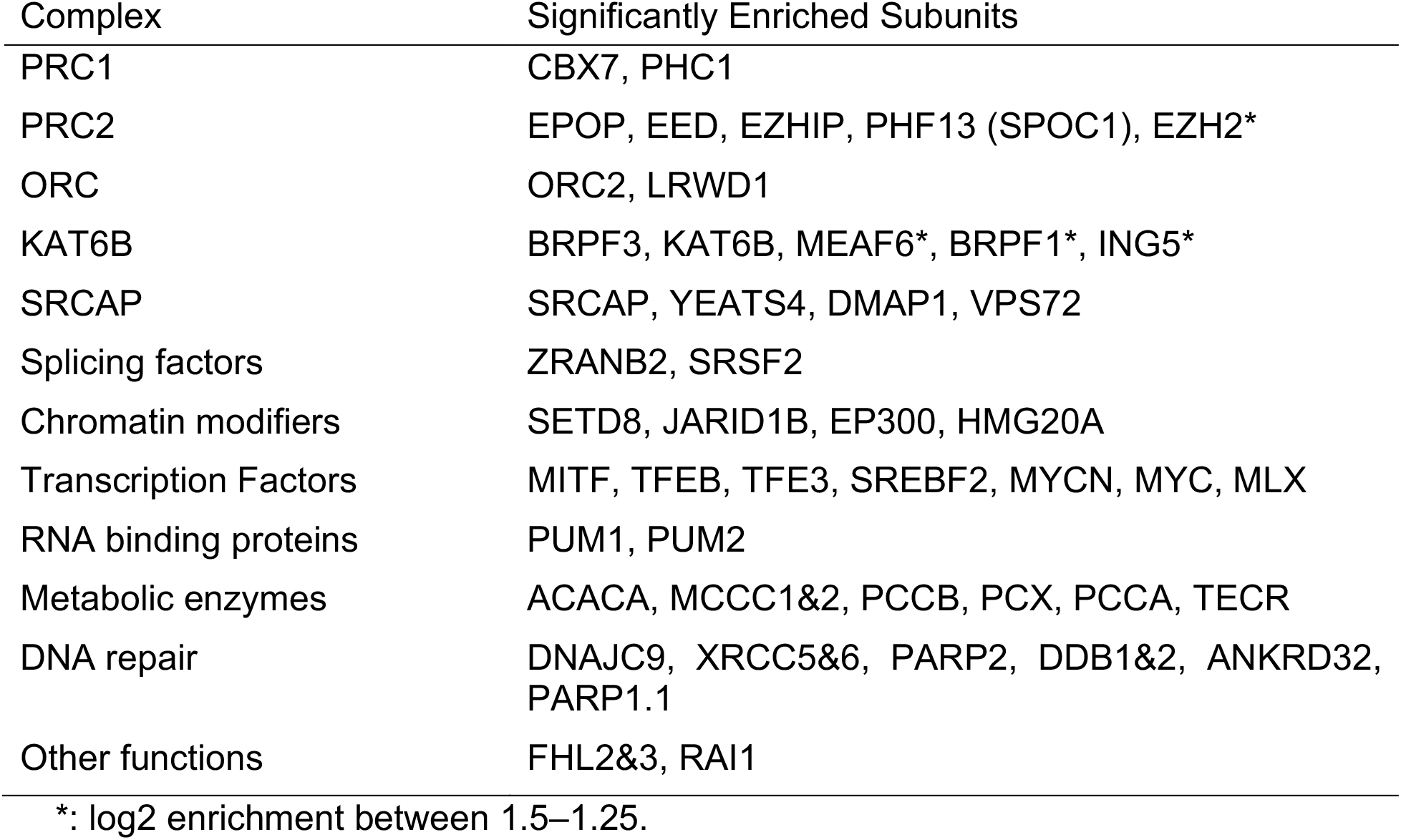
Proteins significantly enriched for asymmetric bivalent nucleosomes.

## Methods

### Cell culture

E14 ESCs were grown on plates coated with gelatin (0.1%, Sigma) in DMEM containing 4.5 g/l glucose (Gibco) supplemented with 15% fetal bovine serum (Gibco, South American, A3160802), 2 mM l-glutamine, 1 mM pyruvate, 1x MEM nonessential amino acids, 50 units/ml penicillin and 50 μg/ml streptomycin (all Gibco), 0.2 mM β-mercaptoethanol (Sigma), and heterologously expressed recombinant LIF (made inhouse and batch-tested for maintenance of self-renewal) at 37°C and 5% CO_2_.

### CRISPR-mediated genome editing of ESCs

To perform genome editing in E14 ES cells, sgRNAs specific to the gene of interest were designed and cloned into pX458 ((addgene #48138)Ran et al., 2013). For homology repair, 200-base single stranded oligodeoxynucleotide (ssODN, IDT) donors were designed to encode the desired mutations or insertions flanked by homology regions of at least 30 bp. Low-passage E14 ESCs were transfected with pX458 plasmid and the corresponding ssODN donor using Lipofectamine 2000 or 3000 (Invitrogen). After GFP fluorescence-based FACS sorting of transfected single cells, single cell colonies were expanded for 7-10 days on 15-cm plates before picking of single colonies, trypsinisation, and further expansion in 96-well plates in duplicate. DNA was extracted from one plate using QuickExtract DNA extraction solution (Epicentre) to perform genotyping by PCR using primer pairs specifically recognizing the mutated sequences introduced by the ssODN and a genomic sequence adjacent to the region of integration. To confirm correct genotypes in cell clones positive for the desired mutations from the genotyping PCR, genomic sequences surrounding and including the integration sites were amplified by PCR and analyzed by Sanger sequencing on PCR-amplified genomic material to confirm presence of the desired mutations only.

### Preparation of ES cell nuclear extract

E14 ESC nuclear extract (NE) was prepared based on modifications of existing protocols (Conaway et al., 1996; LeRoy et al., 1998). E14 ESCs were washed in PBS (Gibco) and harvested by scraping into PBS supplemented with 1 mM EDTA. All subsequent steps were performed at 4°C. Cells were collected by centrifugation at 500 g for 8 min, washed with TMSD (20 mM HEPES pH 7.9, 10 mM KCl, 1.5 mM MgCl_2_, 250 mM sucrose, 0.5 mM DTT, 0.2 mM PMSF), and collected by centrifugation as before. The cells were resuspended in TMSD containing 0.05% NP40 and incubated on ice for 5 min to allow for cell lysis to occur. After successful cell lysis was confirmed by staining an aliquot of the cell suspension with Trypan Blue (Gibco), nuclei were pelleted by centrifugation for 10 min at 1000 g. The supernatant (cytosolic fraction) was removed and the pellet containing nuclei resuspended in BC420 buffer (20 mM HEPES pH 7.9, 420 mM KCl, 20% glycerol, 1.5 mM MgCl_2_, 0.2 mM EDTA, 0.5 mM DTT, 0.2 mM PMSF) and incubated at 4°C with gentle rotation for 1 h. After a 15-s vortex, the suspension was centrifuged for 30 min at 25,000 g. The supernatant was dialysed three times against 50 volumes of BC150 buffer (20 mM HEPES pH 7.9, 150 mM KCl, 20% glycerol, 1.5 mM MgCl_2_, 0.2 mM EDTA, 0.5 mM DTT, 0.2 mM PMSF) for 1 h each. The dialysate was centrifuged for 30 min at 25,000 g. The resultant supernatant is the NE. For all nucleosome pulldowns, 3 independent extractions were performed and the resulting NEs combined to compensate for batch variations between individual preparations.

### Expression and purification of histones

*Xenopus* H3 and H4 and human H2A and H2B were expressed from pET-3a or pET-3d vectors in BL21 (DE3) pLysS for H3, H2A, and H2B or BL21 (DE3) for H4 through induction with 0.2 mM IPTG for 4 h at 37°C. Histones were purified from inclusion bodies and solubilised in unfolding buffer (20 mM Tris-HCl pH 7.5, 7 M guanidine HCl, 10 mM DTT). Extracted histones were dialysed against three changes of urea dialysis buffer (10 mM Tris HCl pH 8, 7 M urea, 100 mM NaCl, 1 mM EDTA, 5 mM β-mercaptoethanol). This and all subsequent histone dialysis steps were carried out at 4°C. Histones were then purified further by passing over a HiTrap Q column (GE Healthcare) before binding to and NaCl gradient elution from HiTrap SP cation exchange chromatography columns (GE Healthcare). Fractions containing histones were pooled and dialyzed three times against water containing 5 mM β-mercaptoethanol and lyophilised for long-term storage at −80°C.

To express histones for native chemical ligation (NCL), constructs encoding truncated *Xenopus* histone H3 were generated in pET-3a. For generation of H3K4me3- and H3K27me3-modified histones, truncated H3 lacking residues 1–31 after the initiator methionine, with a threonine-to-cysteine substitution at position 32 of *Xenopus* H3 and a cysteine-to-alanine substitution at position 110 (H3Δ1–31T32C C110A) was expressed in BL21 (DE3) pLysS and purified as above, except for the final dialysis, which was carried out as two rounds of dialysis against 1 mM DTT in H_2_O and one round against 0.5 mM TCEP (Sigma) before lyophilisation and storage. For generation of H3K27me3-modified histones for some experiments, a similarly truncated Xenopus H3 construct was used, lacking the first 44 residues and carrying a threonine-to-cysteine mutation at residue 45 (H3Δ1–44T45C C110A).

### Generation of modified histones by native chemical ligation

NCL reactions were carried out in 6 M Guanidine HCl, 250 mM sodium phosphate buffer pH 7.2, 150 mM 4-mercaptophenylacetic acid (MPAA, Sigma), 50 mM TCEP for 72 h at room temperature with constant agitation. Reactions were then dialyzed three times against urea dialysis buffer (see above, but with 1 mM DTT instead of 5 mM β-mercaptoethanol). Ligated full-length modified histone was separated from unligated histone through cation exchange chromatography on a HiTrap SP column (GE Healthcare) and then dialysed against three changes of water containing 5 mM β-mercaptoethanol before lyophilization and storage at −a80°C until use. For H3K4me3-or H3K27me3-modified histones, H3Δ1–31T32C C110A was reacted with a synthetic peptide spanning residues 1–31 of histone H3.1 containing trimethylated lysine at position 4 or 27, respectively, and a C-terminal benzyl thioester (Peptide Protein Research). For some H3K27me3-modified histones, H3Δ1–44T45C C110A was reacted with a synthetic peptide spanning residues 1–44 of histone H3.1 including trimethylated lysine at position 27 and a C-terminal benzyl thioester.

### Reconstitution of recombinant nucleosomes

To reconstitute histone octamers, the four core histones were resuspended in unfolding buffer (see above), mixed in a mass ratio of 1:1:1.2:1.2 (H3:H4:H2A:H2B), and dialysed against three changes of refolding buffer (10 mM Tris HCl pH 8, 2 M NaCl, 1 mM EDTA, 5 mM β-mercaptoethanol) at 4°C. After centrifugation to remove precipitate formed during dialysis, correctly assembled histone octamers were purified by size exclusion chromatography in refolding buffer on a S200 column (GE Healthcare), using an Akta PURE system (GE Healthcare).

Asymmetrically modified octamers were further purified as described previously (Voigt et al., 2012), with slight modifications; the two copies of histone H3 incorporated into asymmetric octamers were differentially tagged with either His or Strep tag. Sequential affinity chromatography was performed on the combined fractions containing the desired asymmetric protein complexes from the size exclusion chromatography step by first using a Ni-sepharose HisTrap column (GE Healthcare) and then a StrepTrap HP column (GE Healthcare). Symmetric His-His-tagged complexes and the desired asymmetric complexes containing one His- and one Strep-tagged histone H3 were eluted from the HisTrap column with a gradient (0-250 mM) of imidazole in elution buffer (2 M NaCl, 25 mM Tris pH 8). Fractions containing His-tagged protein as verified by Western blot with an anti-His antibody (Sigma) were pooled and loaded onto the StrepTrap column. Symmetric His-His-tagged complexes eluted in the flowthrough, whereas asymmetric His-Strep-tagged complexes were eluted from the StrepTrap column in elution buffer containing 2.5 mM desthiobiotin (IBA). After each chromatography step, elution profiles and sample purity were confirmed by Western blot with antibodies against His tag (Sigma, H1029, lot 033m4785) and Strep tag (IBA, 2-150-7001).

The DNA template for mononucleosome assembly was generated by PCR with a biotinylated forward primer, amplifying a 209-bp fragment centred around the 147-bp 601 nucleosome positioning sequence (Lowary and Widom, 1998), followed by PCR purification and elution into TE buffer (10 mM Tris pH 8, 1 mM EDTA). To reconstitute recombinant mononucleosomes, DNA and histone octamers were combined in refolding buffer supplemented with 5 M NaCl to compensate for reduction in NaCl concentration due to introduction of TE buffer with the DNA, followed by gradient dialysis against TE buffer down to 400 mM NaCl and then a step dialysis against TE buffer at 4°C. Optimal ratios of DNA and histone octamer were determined so that at least 95% of DNA was complexed while avoiding over-assembly and unspecific DNA binding of histones. Assemblies were routinely checked by native gel electrophoresis on 6% acrylamide gels in TGE buffer.

### Nucleosome pulldown assays

For pulldown assays with recombinant modified nucleosomes and E14 ESC NE, streptavidin sepharose high performance beads (GE Healthcare, 10.5 μl of slurry per pulldown) were briefly washed three times with pulldown buffer (20 mM HEPES pH 7.9, 150 mM NaCl, 10% glycerol, 1 mM EDTA, 1 mM DTT, 0.2 mM PMSF, 0.1% v/v NP-40). All centrifugation steps were carried out at 1,500 g for 2 min at 4°C. All incubation steps were carried out with constant rotation at 30 rpm on a rotator mixer (Starlab) at 4°C in the cold room. After washes, beads were incubated overnight with 10.5 μg of assembled recombinant nucleosomes (in TE buffer diluted with pulldown buffer and adjusted to a final concentration of 0.1% NP-40 and 150 mM NaCl by addition of 10% NP-40 and 5 M NaCl, respectively). For pulldown titrations with increasing amounts of recombinant nucleosomes, nucleosomes were assembled in 4 batches of 10 μg each, combined, and incubated with beads in the amounts indicated in Fig. 1e, g. The amount of beads used was scaled with the amount of nucleosomes (6.4, 6.4, 12, 24 μl slurry for 3, 6, 12, 24 μg of nucleosomes). Beads were then collected by centrifugation and washed briefly with three changes of pulldown buffer. Bead-bound nucleosomes were incubated with 500 μg of NE for 2 h. Beads were then washed with pulldown buffer by 5-min incubations under rotation, first with two washes of pulldown buffer containing NP-40, followed by three washes of pulldown buffer without NP-40.

After washes, bound proteins were eluted from beads. For subsequent LC-MS/MS analysis, beads were resuspended in elution buffer (2 M Urea, 100 mM Tris pH 7.5, 10 mM DTT) and incubated for 20 min in a thermomixer (Eppendorf) at 1000 rpm at 25°C. After incubation, iodoacetamide (Sigma) was added to a final concentration of 55 mM and incubated for 10 min in the thermomixer. The eluted proteins were digested with 0.3 μg of trypsin (Thermo Fisher Scientific) for 2 h whilst shaking and then collected by centrifugation at 1500 g for 2 min. The supernatant containing eluted peptides was removed and 50 μl elution buffer was added to the beads followed by 5 min incubation in a thermomixer. The beads were centrifuged at 1,500 g for 2 min and the two elutions combined. 0.15 μg trypsin added and the solution was digested overnight at room temperature before digestion was stopped by the addition of 10% TFA. For LC-MS/MS LFQ analysis, pulldowns were performed in triplicate to ensure robustness of LFQ-MS analysis.

For Western Blot analysis, elution was performed by boiling for 5 min at 95°C with 1.5x SDS sample buffer (95 mM Tris HCl pH 6.8, 15% glycerol, 3% SDS, 75 mM DTT, 0.15% bromophenol blue). 30% of bound sample was loaded for Western blot analysis. Binding was analyzed by probing with antibodies against TAF3 (abcam, AB188332, lot GR220332-3), ING2 (abcam, AB109594, lot YH102514), PHF2 (clone D45A2, Cell Signaling, 3497s, lot 2), CBX7 (abcam, AB21873, lots GR3210651-1 and −2), EZH2 (BD Biosciences, 612667, lot 415817), RING1B (clone D22F2, Cell Signaling, 5694s, lot 3), HA tag (abcam, AB9110, lot GR3199553-1, or Cell Signaling, clone C29F4, 3724s, lot 9). Antibodies against histone H4 (Cell Signaling, 13919, lot 3), H3K4me3 (abcam, ab8580, lot GR273043-2), and H3K27me3 (Cell Signaling, 9733, lot 8) were used as loading controls. Blots were developed with Clarity ECL reagent (Bio-Rad) and imaged with a Chemidoc Touch imaging system (Bio-Rad). For Western analysis, pulldowns were performed in duplicate or triplicate.

### LC-MS/MS analysis

Following digestion, half of each sample was diluted with an equal volume of 0.1% TFA and loaded onto home-made StageTips as described (Rappsilber et al., 2003). Peptides were eluted in 40 μl 80% acetonitrile in 0.1% TFA and concentrated down to 1 μl by vacuum centrifugation (Concentrator 5301, Eppendorf). Samples were then prepared for LC-MS/MS analysis by diluting them to 6 μl with 0.1% TFA.

LC-MS/MS analyses for samples that were digested on beads were performed on a Q Exactive mass spectrometer and for samples processed by in-gel digestion on an Orbitrap Fusion Lumos Tribrid mass spectrometer (both Thermo Fisher Scientific), both coupled on-line to Ultimate 3000 RSLCnano Systems (Dionex, Thermo Fisher Scientific). Peptides were separated on a 50-cm EASY-Spray column (Thermo Fisher Scientific) assembled in an EASY-Spray source (Thermo Fisher Scientific) and operated at 50°C. In both cases, mobile phase A consisted of 0.1% formic acid in water, while mobile phase B consisted of 80% acetonitrile and 0.1% formic acid in water. Peptides were loaded onto the column at a flow rate of 0.3 μl/min and eluted at a flow rate of 0.25 μl/min with the following gradient: 2-40% buffer B in 150 min, then to 95% in 11 min. For Q Exactive runs, FTMS spectra were recorded at 70,000 resolution (scan range 350-1400 m/z) and the ten most intense peaks of the MS scan with charge ≥ 2 were selected with an isolation window of 2.0 Thomson for MS2 (filling 1.0E6 ions for MS scan, 5.0E4 ions for MS2, maximum fill time 60 ms, dynamic exclusion for 50 s). For Orbitrap Fusion Lumos runs, survey scans were performed at 120,000 resolution (scan range 350-1400 m/z) with an ion target of 4.0e5. MS2 was performed in the ion trap at a rapid scan rate with ion target of 2.0E4 and HCD fragmentation (Olsen et al., 2007) with a normalized collision energy of 27. The isolation window in the quadrupole was 1.4. Only ions with a charge between 2 and 7 were selected for MS2.

### Processing and visualisation of LC-MS/MS data

The MaxQuant software platform (Cox and Mann, 2008) version 1.6.1.0 was used to process raw files and searches were conducted against the Mus musculus complete/reference proteome (Uniprot, released in July, 2017), using the Andromeda search engine (Cox et al., 2011). The first search peptide tolerance was set to 20 ppm while the main search peptide tolerance was set to 4.5 ppm. Isotope mass tolerance was 2 ppm and maximum charge was set to 7. A maximum of two missed cleavages was allowed. Carbamidomethylation of cysteine was set as fixed modification. Oxidation of methionine and acetylation of the N terminus were set as variable modifications. Label-free quantification (LFQ) analysis was performed by employing the MaxLFQ algorithm as described (Cox et al., 2014). Peptide and protein identifications were filtered to 1% FDR.

Data was analysed and visualised in R version 3.5.3. The DEP package (Zhang et al., 2018) version 1.4.1 was used to determine differential enrichment of proteins between nucleosome pulldowns. Data was filtered on reverse hits and potential contaminants, before missing values were filtered to retain only proteins quantified in at least two replicates across all nucleosome pulldown conditions and replicates analysed. The data was normalized using vsn, and imputation was carried out using the MinProb function in DEP. Proteins were considered differentially enriched when the log2 fold change was greater than 1.5, and the p value was lower than 0.01. Bar plots of fold changes of LFQ intensities relative to the mean LFQ intensity for select proteins were generated in DEP using the accompanying Shiny package. Volcano plots were generated in ggplot2.

### Chromatin immunoprecipitation

To prepare chromatin for ChIP, E14 ESCs were grown as described above. Crosslinking was carried out directly on 15-cm culture plates with 1% formaldehyde in fixation buffer (DMEM supplemented with 10 mM HEPES pH 7.6, 15 mM NaCl, 0.15 mM EDTA, 0.075 mM EGTA for 10 min at room temperature with gentle rocking. Crosslinking was stopped by addition of glycine to 0.125 M final concentration and incubation for 3 min at room temperature with gentle rocking. Fixed cells were washed with cold PBS, scraped off the plate in cold PBS, and collected by centrifugation at 2500 g for 5 min at 4°C. Cell pellets snap frozen in liquid nitrogen and stored at −80°C. To prepare fragmented chromatin, cell pellets were resuspended in lysis buffer 1 (50 mM HEPES pH 7.6, 140 mM NaCl, 1 mM EDTA, 10% glycerol, 0.5% NP-40, 0.25% Triton-X100) and incubated with constant rotation for 10 min at 4°C. Cells were then centrifuged at 3000 g for 5 min and cell pellets resuspended in lysis buffer 2 (10 mM Tris-HCl pH 8.0, 200 mM NaCl, 1 mM EDTA, 0.5 mM EGTA) and incubated and centrifuged as for lysis buffer 1. Pellets were then resuspended in lysis buffer 3 (10 mM Tris-HCl pH 8.0, 1 mM EDTA, 0.5 mM EGTA, 0.5% N-Lauryl Sarcosine) at a density of 75 mg cell pellet weight/ml. The suspension was transferred to polystyrene tubes (BD Falcon) and sonicated using a Bioruptor (Diagenode) at maximum output setting using 30 s ON, 30 s OFF cycles. Number of cycles (15-20) was experimentally determined to yield decrosslinked fragments of 250-350 bp size. Size of chromatin fragments was verified on a 1% agarose gel and on a high sensitivity DNA chip in a 2100 bioanalyzer (Agilent) after decrosslinking overnight at 65°C with shaking in elution buffer (100 mM NaHCO3, 1% SDS, 200 mM NaCl, 0.4 mg/ml each proteinase K and RNAse A). Sonicated suspensions were clarified by centrifugation at 20,000 g for 10 min at 4°C. DNA concentration as a proxy for chromatin concentration was determined by Nanodrop ND-1000 (Thermo Fisher Scientific).

For ChIP, chromatin was clarified by centrifugation at 20,000 g for 10 min and incubated overnight with antibodies in 1x IP buffer (10 mM Tris-HCl pH 8.0, 150 mM NaCl, 1 mM EDTA, 0.5 mM EGTA, 0.25% N-Lauryl Sarcosine, 1% Triton X100) with constant rotation at 4°C. Per IP, 50 μl protein A or G magnetic dynabeads (Invitrogen) were prepared by washing twice, then incubating overnight with PBS containing 0.5% (w/v) BSA overnight at 4°C with rotation, then washing 5 times in TE buffer (10 mM Tris pH 8, 1 mM EDTA), and finally resuspending in TE. After overnight incubation, IPs were centrifuged at 20,000 g for 10 min at 4°C and the supernatants were combined with blocked dynabeads and incubated at 4°C for 2-3 h under constant rotation. Beads were then washed 5 times in cold RIPA buffer (10 mM HEPES pH 7.6, 1 mM EDTA, 0.5 M LiCl, 1% NP-40, 0.1% N-Lauryl Sarcosine) with 2 min rotating incubation at room temperature for each wash. Beads were then washed briefly with TEN buffer (TE with 50 mM NaCl). Immunoprecipitated DNA was eluted from washed beads by addition of elution buffer (100 mM NaHCO3, 1% SDS, 200 mM NaCl, 0.4 mg/ml each proteinase K and RNAse A) and incubation overnight at 65°C with constant agitation to reverse crosslinking. The next day samples were purified using PCR cleanup kit columns (Monarch, NEB) and eluted using 50 μl kit elution buffer.

Suitable dilutions of eluted DNA were used for qPCR analysis on a Lightcycler 480 (Roche) using qPCRBIO SyGreen Blue Mix (PCRBiosystems). Enrichments were calculated as percentage of input. The following primers were used for qPCR analysis: Polm, tgacgggcacaattacacca, aaaggcttccgcgtcctaga; Gapdh, ggg ttc cta taa ata cgg act gc, ctg gca ctg cac aag aag atg; Pou5f1 (Oct4), ggc tct cca gag gat ggc tga g, tcg gat gcc cca tcg ca; Nanog, TGGCCTTCAGATAGGCTGAT, caagaagtcagaaggaagtgagc; HoxC5, gtactgctacggcggattgg, taccccgtggagagagttgg; Olig1, gggttacaggcagccaccta, atgcggtggaagaggatgag; Fgf5, GCGACGTTTTCTTCGTCTTC, ACGAAACCCTACCGGACTCT; HoxA7, GAGAGGTGGGCAAAGAGTGG, CCGACAACCTCATACCTATTCCTG; Fabp7, TGAGCAAATCACAAGGAGGA, TGGAGGAACTCGGGTCTTAC; Bcor promoter, gtaaaaccgaaagcgagcaa, GAGGGTTTCTCCTCCGACTT; Bcor gene body, GGGGGTAACTGTGGGAATCT, TCTCCACACCTTCAGCCTCT; Fzd1, ACATGAGCCCGTAAACCTTG, GGTGCCCTCCTACCTCAACT.

The following antibodies were used for ChIP: H3K4me3 (Cell Signaling, 9751, lot 10, 4 μg antibody for 40 μg chromatin), H3K27me3 (Cell Signaling, 9733, lot 14, 4 μg antibody for 40 μg chromatin), H3K23ac (abcam, AB177275, lot GR3213108-2, 2 μl antibody for 40 μg chromatin), H3K14ac (abcam, AB52946, lot GR3252548-2, 4 μl antibody for 40 μg chromatin), RING1B (Cell Signaling, 5694, lot 3, 4.5 μl antibody for 50 μg chromatin), and HA tag (Cell Signaling, 3724, lot 9, 4 μl antibody for 200 μg chromatin).

### Bioinformatic analysis of ChIP-seq data

Obtained sequences were filtered for low-quality reads, and adapter sequences were removed using Trim Galore. Filtered reads were mapped to the mouse genome (version mm9) using the BOWTIE in QuasR (Gaidatzis et al., 2015) using default settings. Identical reads from PCR duplicates were filtered out from the obtained .bam files using Samtools rmdup. Wiggle tracks, for visualisation in the UCSC genome browser, were obtained using QuasR qExportWig() on the aligned reads. Heatmaps and average density-profiles 4 kb around RefSeq TSS were generated from .bam files using genomation in R using a 500 bin approach and strand-aware settings (Akalin et al., 2015). TSS were sorted into three categories based on the obtained binding profiles. Towards this, the k-means clustering option was applied using H3K4me3, H3K27me3, TAF3, and CBX7 profiles to calculate the clusters. The three clusters representing TSS in bivalent state, H3K4me3-only TSS, and TSS with low signal were then used to calculate average density around TSS.

### Western Blot Analysis of ESC whole-cell extracts

To generate ESC whole cell extract for Western Blot analysis, ESCs were washed with PBS and detached from culture plates by trypsinisation (Gibco). After addition of ESC culture media to stop trypsin digest, cells were collected, resuspended, and counted using a haemocytometer. Suitable amounts of ESCs were collected by centrifugation at 400 g for 5 min at room temperature and lysed by resuspending in a volume of 1X SDS sample buffer (63 mM Tris HCl pH 6.8, 10% glycerol, 2% SDS, 50 mM DTT, 0.1% bromophenol blue) suitable for loading onto an SDS-PAGE gel. For histone modification analysis, the equivalent of 3×10^4 cells were loaded per lane. For chromatin factor analysis, 5-10×10^5 cells were used per lane for Western Blot analysis.

Expression or modification status was analyzed by probing with antibodies against EZH2 (BD Biosciences, 612667, lot 415817), HA tag (Cell Signaling, clone C29F4, 3724s, lot 9), KAT6B/MORF (abcam, ab191994, lot GR3180184), H3K4me3 (abcam, ab8580, lot GR273043-2), H3K23ac (abcam ab177275, lot GR3213108-2), and H3K27me3 (Cell Signaling, 9733, lot 8). Antibody against histone H4 (Cell Signaling, 13919, lot 3) was used as a loading control. Blots were developed with Clarity ECL reagent (Bio-Rad) and imaged with a Chemidoc Touch imaging system (Bio-Rad). Western analysis was performed in duplicate.

### Purification of endogenously FLAG-HA-tagged KAT6B from E14 ESCs

About 1.625 x 10^9 ESCs expressing endogenously FLAG-HA-tagged KAT6B were grown as described above for E14 ESCs. NE was generated as described above, albeit using 0.1% NP40 in the TMSD lysis step. After solubilization in BC420, vortexing, and centrifugation as above, the resulting NE was adjusted to 300 mM KCl using BC buffer without KCl (BC0) by monitoring conductivity with a conductivity meter (Mettler Toledo). To prepare Anti-FLAG M2 affinity agarose beads (Sigma) for affinity purification, beads were incubated with 100 mM glycine, washed with 1M HEPES pH 7.9, and washed to equilibrate with BC300 buffer (see BC420 above but with 300 mM KCl). The pretreated and washed anti-FLAG beads were combined with the NE and incubated for 2 h with constant rotation on a rotator mixer (Starlab) at 4°C in the cold room. After incubation, beads were collected by centrifugation at 800 g for 2 min at 4°C and washed twice with BC300 buffer and then once with BC100 buffer (see BC420 above but with 100 mM KCl). Bound FLAG-HA-tagged KAT6B and associated proteins were eluted with 0.3 μg/μl 3x FLAG tag peptide (Sigma) in BC100 buffer for 1 h with gentle rotation at 4°C.

### Histone acetyltransferase assays

Recombinant nucleosomes were reconstituted as described above using a plasmid containing 12 177-bp repeats containing the 147-bp 601 sequence. Nucleosomes and KAT6B enzyme preparations were combined in HAT buffer (50 mM Tris-Cl pH 7.5, 10% glycerol, 0.1 mM EDTA) and 4 mM DTT. Reactions were started by the addition of ^3^H-acetyl coA (90 kBq; Hartmann) and incubated at 30°C for 2 h. Reactions were stopped addition of 3x SDS sample buffer (see above) to a final concentration of 1x by boiling for 5 min at 95°C with. Proteins were separated by SDS-PAGE, transferred to PVDF membrane, and stained with Coomassie to assess loading. Incorporated ^3^H-acetyl was detected using BioMax MS film (Kodak, Sigma) and BioMax Transcreen LE amplifying screens (Kodak, via Sigma).

### LC-MS/MS analysis of HA-tagged KAT6B complex purified from E14 ESC

4x NuPAGE LDS Sample buffer (Invitrogen) and 1 M DTT was added to purified endogenously FLAG-HA-tagged KAT6B complexes to a final concentration of 1x and 50 mM, respectively. Samples were boiled at 95°C for 5 min before running on a NuPAGE 4-12% Bis-Tris protein gradient mini gel (Invitrogen) at 200 V for 5 min in MOPS running buffer (Invitrogen) to concentrate the sample. Gels were washed three times with deionized water for 5 min each before staining with InstantBlue (Expedeon) for 1 h. Stained gels were washed three times with deionized water for 5 min each to remove excess stain. For in-gel digestion, the stained band containing the protein sample was excised and de-stained by washing with 50 mM ammonium bicarbonate (Sigma) and 100% (v/v) acetonitrile (Sigma). Proteins were reduced in 50 mM ammonium bicarbonate with 10 mM DTT for 30 min at 37°C and alkylated in 55 mM iodoacetamide (Sigma) for 20 min at ambient temperature in the dark. Proteins were then digested overnight at 37°C with 12.5 ng/μl trypsin (Pierce). Samples were then analyzed by LC-MS/MS as described above.

### Neuronal differentiation of ESC

Neuronal differentiation was performed as previously described (Bibel et al., 2007). Embryoid body (EB) formation was induced by removal of LIF. On day 0, ESCs were plated onto non-adherent bacterial plates (Greiner) in 10 ml EB medium (ESC medium described above but without LIF and with FBS reduced to 10%) at a density of 4 x 10^6 cells per 10-cm plate. On day 2, the media was changed by gently transferring EBs to a 15-ml conical tube (Sarstedt), allowing EBs to settle for 5 min, then carefully aspirating the supernatant. EBs were then gently resuspended in 13 ml of fresh EB media using a 5-ml serological pipette (Sarstedt) and transferred to a fresh plate. On day 4 and day 6, media was changed in the same manner to 15 ml EB medium with 10 μM all-trans retinoic acid (Sigma). On day 8, EBs were washed 3 x in 20 ml PBS, trypsinised (Sigma, prepared at 0.5 mg/ml in 0.05% v/v EDTA/PBS) for approximately 3 min at 37°C with agitation and then quenched in 10 ml EB medium. Dissociated neural progenitor cells were then centrifuged for 5 min at 300 g, resuspended in 10 ml EB medium and passed through a 40-μm cell strainer (Thermo Fisher Scientific). Cells were counted, centrifuged again and resuspended in Advanced DMEM/F12 (Gibco) with 1x N2 supplement (Gibco) before being plated onto poly-D-lysine (Sigma) and laminin (Sigma) coated 6-cm dishes at a density of 3 x 10^6 cells/plate. On day 9, half of the media was replaced with Neurobasal medium (Gibco) containing 1x B27 supplement (Gibco). This was repeated on day 12.

### RNA preparation and RT-qPCR

Cells were resuspended in 1ml TriPure isolation reagent (Roche) per 10-cm plate of ESCs or EBs (or per 2x 6-cm plate for plated neural cells) and RNA extracted following the manufacturer’s instructions, resuspended in BTE (10 mM Bis-Tris pH 6.7, 1 mM EDTA), and treated with Turbo DNAse (Ambion) for 30 min at 37°C. 1 ml TriPure was then added and RNA extraction repeated before resuspending the RNA pellet in 50 μl BTE. Integrity of RNA was assessed using RNA 6000 Nano chips on a Bioanalyzer (Agilent). cDNA was prepared from of 1 μg RNA per sample using SuperScript IV reverse transcriptase (Invitrogen). qRT-PCR experiments with the resulting cDNA samples were carried out on a Lightycler 480 (Roche) using qPCRBIO SyGreen Blue Mix (PCRBiosystems). Results were analysed using the 2-ΔΔCT method with Gapdh as the reference.

Primers used for qPCR were: GAPDH, CATGGCCTTCCGTGTTCCT, GCGGCACGTCAGATCCA; Oct4, AGATCACTCACATCGCCAATCA, CGCCGGTTACAGAACCATACTC; Otx2, CATGATGTCTTATCTAAAGCAACCG, GTCGAGCTGTGCCCTAGTA; Nanog, AGGCTTTGGAGACAGTGAGGTG, TGGGTAAGGGTGTTCAAGCACT; Fgf5, AAACTCCATGCAAGTGCCAAAT, TCTCGGCCTGTCTTTTCAGTTC; Mash1, TAACTCCCAACCACTAACAGGC, TGAGGAAAGACATCAACCCAG; HoxB13, CATTCTGGAAAGCAGCGTTTG, TGTTGGCTGCATACTCCCG; Pax6, GATTGAGGCTCTGGAGAAAG, CCATTTGGCCCTTCGATTAG; Fabp7, TGTAAGTCTGTGGTTCGGTTG, AGCAACGATATCCCCAAAGG; Eed, CAGCCACCCTCTATTAGCAG, GCATTTCCATGGCCAACATAG; Polm, ATGGGCTGTTTGATCCTGAG, ACAGGCATTTCTCAGTTCAGG; Tbp, TGTATCTACCGTGAATCTTGGC, CCAGAACTGAAAATCAACGCAG; Polr2a, GGTCCTTCGAATCCGCATC, CAGGGTCATATCTGTCAGCATG; Sox2, GGGAGAAAGAAGAGGAGAGAGA, GCCGCGATTGTTGTGATTAG; Tbx3, AGGAGCGTGTCTGTCAGGTT, GCCATTACCTCCCCAATTTT; Ring1b, CGAACACCTCAGGAGGCAATA, ATACAATCCGCGCAAAACCG; Kmt2b, AAGTTCCGCTACCACCAGC, TGTCAAAGGTACACTTCCGGAGATA.

### Immunofluorescence analysis

Cells were grown on 16-mm glass coverslips in 12-well plates. Coverslips were coated with 0.1% gelatin (ESCs) or with poly-D-lysine and laminin (for neural cells). Cells were fixed by washing once in PBS and adding 4% paraformaldehyde (Sigma) for 20 min at room temperature. Coverslips were then washed 3 x 10 min in PBS then stored in PBS at 4°C. To carry out immunofluorescence analysis, fixed cells were washed once in PBS and blocked in 10% donkey serum (Sigma) in PBS with 0.1% Triton X-100 for 1 h and incubated overnight with primary antibodies at 4°C. The following day, cells were incubated with Alexa Fluor 488 or 555 secondary antibodies (Invitrogen) at a dilution of 1:1000 in 1% donkey serum in PBS with 0.1% Triton X-100 for 1 h at room temperature in the dark. Coverslips were then washed 3x with PBS for 10 min. Nuclei were counterstained with 50 nM DAPI (Roche) for 5 min, and washed 2x 10 min with PBS before being mounted on slides using vectashield (Vector Laboratories). Imaging was carried out using a Zeiss Axio imager with a 40x objective. The microscope was equipped with a Hamamatsu Flash sCMOS camera. Micromanager software (version 1.4) was used to capture images (Edelstein et al., 2014).

